# SC2Spa: a deep learning based approach to map transcriptome to spatial origins at cellular resolution

**DOI:** 10.1101/2023.08.22.554277

**Authors:** Linbu Liao, Esha Madan, António M. Palma, Hyobin Kim, Amit Kumar, Praveen Bhoopathi, Robert Winn, Jose Trevino, Paul Fisher, Cord Herbert Brakebusch, Gahyun Kim, Junil Kim, Rajan Gogna, Kyoung Jae Won

**Affiliations:** Biotech Research and Innovation Centre (BRIC), University of Copenhagen, Ole Maaløes Vej 5, 2200 Copenhagen N, Denmark; Department of Surgery, Virginia Commonwealth University, School of Medicine, 1200 E Broad St. P.O. Box 980011, Richmond VA, 23298, USA; Massey Cancer Center, Virginia Commonwealth University, Richmond, VA, 23298, USA; Institute of Molecular Medicine, Department of Human and Molecular Genetics, Virginia Commonwealth University, School of Medicine, Richmond, VA, 23298, USA; Department of Human and Molecular Genetics, Virginia Commonwealth University, School of Medicine, Richmond, VA, USA; Instituto Superior Tecnico, Universidade de Lisboa, 1049-001 Lisboa, Portugal; Department of Bioinformatics, Soongsil University, 369 Sangdo-Ro, Dongjak-Gu, Seoul 06978, Republic of Korea; School of Systems Biomedical Science, Soongsil University, 369 Sangdo-Ro, Dongjak-Gu, Seoul 06978, Republic of Korea; Department of Computational Biomedicine, Cedars-Sinai Medical Center, Los Angeles, CA, USA

**Author notes:** To whom correspondence should be addressed. RG (Tel: +1□) and KJW.

**Keywords:** spatial transcriptomics, spatial inference, spatial mapping, deep learning, spatially variable genes

## Abstract

**Background:** Understanding cellular heterogeneity within tissues hinges on knowledge of their spatial context. However, it is still challenging to accurately map cells to their spatial coordinates.

**Results:** We present SC2Spa, a deep learning-based approach that learns intricate spatial relationships from spatial transcriptomics (ST) data. Benchmarking tests show that SC2Spa outperformed other predictors and accurately detected tissue architecture from transcriptome. SC2Spa successfully mapped single cell RNA sequencing (scRNA-seq) to Visium assay, providing an approach to enhance the resolution for low resolution ST data. Our test showed that SC2Spa performs well for various ST technologies and robust to spatial resolution. In addition, SC2Spa can suggest spatially variable genes that cannot be identified from previous approaches.

**Conclusions:** SC2Spa is a robust and accurate approach to provide single cells with their spatial location and identify spatially meaningful genes.

## Background

Cells function within the context of tissue architecture and continuously interact with their local environment. The transcriptome of a cell varies in accordance with the location and the environment. For instance, cells express genes related to cell communication (such as ligands and receptors) and genes involved in downstream pathways [1]. Interaction with the neighboring cells influences the transcriptome of a cell [2–6]. Also, there are genes whose spatial expression patterns are associated with tissue architecture [7]. In developing drosophila embryos, several genes showed gradual pattern along body [8]. These findings demonstrate the influence of global tissue structure on cell transcription.

Single-cell RNA sequencing (scRNA-seq) portrays a snapshot of the gene expression of a cell. This expression profile has been used to identify cell types and dynamic transcriptomic changes [9–12]. However, loss of spatial location makes it hard to interpret scRNA-seq with respect to the influence of cellular neighborhood and spatial location within a tissue. Interestingly, even though the physical connection is severed during scRNA-seq, some studies suggest these isolated cells might still hold clues about their microenvironment [3].

Considering the rich information of cells contained in the transcriptome of cells, can the transcriptome of cells be used to predict their location within tissue and infer tissue architecture? Several systematic approaches have tried to predict spatial origin from the transcriptome [13–20]. Many algorithms have relied on transcriptomic similarities to map a cell to its potential spatial origins [13, 15, 21]. Seurat used mixture models and fitted their parameters by estimating the posterior probability to maximize transcriptomic similarity [14]. SpaOTsc applied the optimal transportation algorithm for spatial reconstruction, followed by cell-cell communication inference using ligand-receptor co-expression [17]. NovoSpaRc reconstructed spatial map assuming continuity in gene expression [16, 20]. Despite these efforts, it is still arduous to map single cells to their spatial origins at a single cell resolution. Previous spatial inference tools lack the ability to generalize over large datasets. In addition, the design of previous studies neglected the relative position between spatial voxels, resulting in the loss of information. The algorithms based on transcriptomic similarity and continuity cannot promise accurate detection of cell location at cellular resolution.

To overcome these challenges, we suggest using deep learning as a method to learn the rules to map cells to their respective location within a tissue. Deep learning has the power to represent complex high-dimensional relationships by combining multiple layers of nonlinear functions [22]. There were attempts to use deep learning in predicting location from transcriptome. DEEPsc was developed to map single cells to spatial voxels based on the transcriptome using deep learning. DEEPsc defined mapping as a binary classification task [19], resulting in modest performance. Tangram applied deep learning to map scRNA-seq to reference ST data [18]. To achieve this, Tangram generates an initial random spatial placement of scRNA-seq then it re-aligns cells in the spatial dimension to maximize the total spatial correlation. DEEPsc and Tangram [18, 19] did not train their network to predict the cell locations directly.

To capture the rules of location mapping from the transcriptome to the location in the weight space of the deep neural networks, we developed SC2Spa. SC2Spa consists of a fully connected neural network (FCNN) designed to predict the location of a cell from its transcriptome. SC2Spa uses ST data (both image based and sequence based) to learn the absolute spatial coordinate (output) from transcriptome (input). The trained model can be used to predict the original location of transcriptome (e.g. scRNA-seq). This strategy is similar to CeLEry [23] which is based on a 3-layered FCNN with augmented learning. Compared with CeLEry, SC2Spa is equipped with strategies to effectively learn the data by providing penalties to the weights that do not contribute to the prediction with deeper layered structure (>6). Besides, SC2Spa is equipped with a strategy that effectively mitigates batch effect when integrating scRNA-seq by applying the Wasserstein Distance [24].

Benchmarking tests on cellular resolution ST data show that SC2Spa outperformed other algorithms in predicting approximate location of cells and recapitulated the original tissue structures from independent ST data and scRNA-seq data. SC2Spa can be applied to various ST technologies and is robust to spatial resolution. SC2Spa trained on low-resolution Visium [25] data still successfully recovered the spatial location of key marker genes from scRNA-seq. Based on the trained weights, SC2Spa identified spatially variable genes (SVGs). SC2Spa is a tool that provides accurate spatial location to scRNA-seq and subsequent analysis.

## Methods

### Data processing for SC2Spa

To normalize the ST data, the unique molecular identifier (UMI) count per cell is normalized to 10,000 and then logarithmically transformed. SC2Spa can choose either polar or Cartesian coordinates. Polar coordinates are useful for circular ST data. The spatial coordinates of the beads were normalized between 0 and 1 using min-max normalization. The FCNN is trained to predict the normalized coordinates of beads.

### Training neural network

SC2Spa is composed of 8 layers (input, output, and 6 hidden layers). The number of nodes for the input layer is the number of genes and the number of the output node is 2 for the spatial coordinates. By default, the number of nodes in these layers is set to 4096, 1024, 256, 64, 16, and 4, respectively. The ReLU function is used for the first five hidden layers.

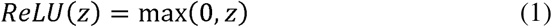

The output layer uses the sigmoid function:

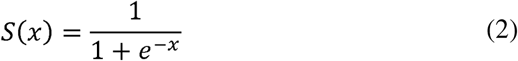

For training, SC2Spa uses the Adam algorithm [26]. The RMSE is the loss function used to train the FCNN. L1 regularization was used to penalize dense layers.

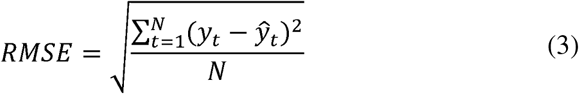

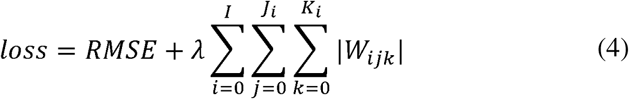

where

*N*: Number of voxels/cells from the reference ST data.

*y_t_*: Spatial coordinates in the reference ST *t*.

*ŷ_t_*: Predicted spatial location on the reference ST *t*

□: coefficient used to adjust the L1 penalty

*W_ijk_*: The *k*-th value in the *j*-th row of the *i*-th weight matrix

*I*: Number of weight matrices

*J_i_*: Number of rows in the *i*-th matrix

*K_i_*: Number of columns of the *i*-th matrix.

To handle the batch effect, Wasserstein distance [24] was used to obtain genes with comparable distributions between them:

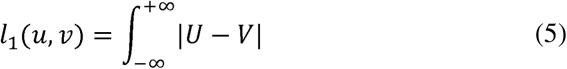

where

*u*: The expression values of a gene in the reference data

*v*: The expression values of a gene in the query data

*U*: The cumulative distribution function (CDF) of *u V*: The CDF of *v*.

Especially for the spatial inference of Visium data [27–29], genes with Wasserstein distances less than 0.1 were selected for training and subsequent analysis to reduce noise associated with low resolution Visium spots.

Glorot normalization [30] is used to initialize the parameters of the neural network. Learning rate 0.001 of Keras [31] is used. Dropout rate is set to 0.05 for the input and the first 5 hidden layers. A dropout layer is used to arbitrarily eliminate a certain number of input nodes during prediction. The batch size is set to 16 for the mouse olfactory bulb dataset, 128 for the other Visium data and 4096 for the other data. Number of epochs is set to 500. The L1 regularization scale factor is set to 0.00000001. Learning rate reduction is configured with a minimal learning rate of 0.00001, reduction factor of 0.5 and patient of 20. Early stopping is configured with a patience parameter of 50. The patience parameter refers to the max number of epochs. These parameters can also be provided by a user.

### Benchmarking

We compared SC2Spa with LR, Novosparc [16, 20], SpaOTsc [17] Tangram [18] and CeLEry [23] in predicting the coordinates of voxels using 5-fold cross validation. While many methods exist for cell-to-spot mapping, most of them only support decomposition at the cell-type level (decomposition) rather than at the single-cell level. Therefore, we specifically selected four methods capable of performing spatial decomposition at single-cell resolution for comparison in our study. Predicted and original coordinates of the ST spots in the leave-out set were used for calculating metrics. LR was performed using Scikit-learn 0.23.1. All parameters of LR were set with default values. pairwise Pearson’s *r* and pairwise RMSE were used to assess the performance. Pairwise refers to all possible cell pairs of the leave-out spots/cells of the 5-fold cross validation tests. The RMSE values were calculated using min-max normalized coordinates. NovoSpaRc [16, 20], SpaOTsc [17] and Tangram [18] were initially designed to find the optimal mapping probability matrix between single cells to spatial voxels. For the 5-fold cross-validation. The leave-out voxels were used as pseudo scRNA-seq data. The remaining data served as the ST reference. The question was reformulated as searching for the optimal transport between the leave-out data and the training data. To optimize SpaOTsc, NovaSparc, and Tangram, we compared two approaches for predicting spatial coordinates: using the coordinates of the most probable spot or employing a probability-weighted sum of spot coordinates. Our benchmark results indicated that SpaOTsc performs best with the former method, while NovaSparc and Tangram achieve superior results with the latter.

When we apply SpaOTsc using all genes, SpaOTsc generated results similar to random mapping. Therefore, we selected the top 1,800 highly variable genes identified using SCANPY [32] for SpaOTsc for all tests except for the drosophila embryo FISH data [15]. For the drosophila embryo FISH data, we used all 84 genes. For the Stereo-seq mouse embryo data, due to the limitation of Random Access Memory (RAM) and the heavy computation cost of SpaOTsc, 55,000 cells out of 74,342 cells were randomly selected and mapped to the 9300 cells in the leave-out set.

### Fine mapping and ST bead reconstruction for spatial origin prediction for scRNA-seq

SC2Spa predicts the spatial coordinates from the transcriptome. The predicted position may not correspond to the center of a bead and multiple single cells can be mapped to a bead. For visualization, the reconstructed expression levels of a bead are the weighted average of the scRNA-seq mapped to the bead. The weight is determined based on the distance from the center of a bead to the predicted cell location. Only the cells within a certain distance to the center of a bead are selected for the ST bead expression reconstruction. We set the distance cutoff as 15µm for Slide-seqV2 and 110µm for Visium.

The weight of a cell *j* for a bead is defined as:

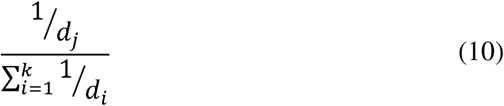

*k*: Number of single cells within a range. The range is the radius of the bead for the ST technology

*d_i_*: Distance between cell *i* and the bead

*d_j_*: Distance between cell *j* and the bead *(j = 1,2,…k)*.

### Quantify the consistency of the prediction with the reference ST data

To evaluate the quality of the prediction, we computed BVI [33, 34] for the gene expression levels between the reconstructed and the reference ST data. The BVI is defined as below:

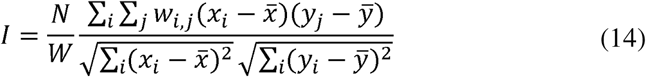

where

*N*: Number of cells

*x_i_*: The expression value of the gene *x* in cell i of the original ST data

*x̄* : Mean of gene *x* expression value in all cells of the reference ST data

*y_i_*: The expression value of the gene *x* in cell *j* of the predicted ST data

*ȳ*: Mean of gene *x* expression value in all cells of the predicted ST data

*w_i,j_*: Weight of a neighboring cell pair cell *i* and cell *j W*: Sum of all *w_ij_*.

The weight of cell pairs with a distance less than a specified threshold is set to 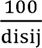 , where “disij“ is the distance between a cell *i* and a cell *j*. 100 is a scale factor. The weight of the remaining pairs of cells is assigned to zero.

### Prioritizing location predictive genes

SC2Spa identifies genes that influence spatial location predictions by analyzing the trained neural network’s weights. It starts at the output layer and traces back through the network, selecting nodes with high weights (top 50% by default) until reaching the input layer, where each node represents a gene. Gene importance is determined by summing the weights connecting it to the selected nodes. This process highlights genes whose expression patterns vary spatially.

An importance score is assigned to each gene by aggregating weight matrices of the NN backward.

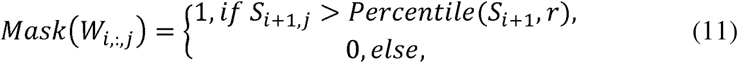

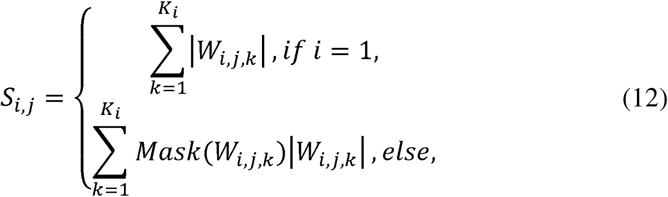

, where

*W_i,j,k_*: The *k*-th value in the *j*-th row of the *i*-th weight matrix

*W_i,:,j_*: The *j*-th column of the *i*-th weight matrix

*S_i,j_*:The sum of the selected values of the *j*-th row of the *i*-th weight matrix

*Percentile*(*S_i+1_,r*): The r-th percentile of S_i+1_. r was set as 50 in our study

I: Number of weight matrices, which is 1 more than the number of hidden layers

*K_i_*: Number of columns of the *i*-th weight matrix.

The calculation was performed backwards from the last weight matrix to the first weight matrix. The output from the first weight matrix is *S_1_*. *S_1_* assesses the significance of genes in determining the spatial location of beads. The importance score of the *j*-th gene *S_1j_* will be normalized subsequently:

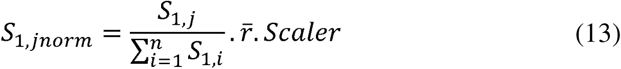

where

*S_1,jnorm_*: The normalized importance score of the j-th gene

*S_1,i_*:The importance score of the *i*-th gene

*r̄*,: The average Pearson correlation coefficient between the prediction and reference spatial coordinates

*Scaler*: A factor for scaling the importance score. Default 1000.

### Prioritizing spatially variable genes

Moran’s I of each gene was calculated using the following equation:

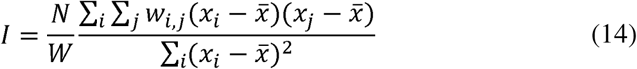

where

*N*: Number of cells

*x_i_*: The expression value of the gene *x* in cell *i*

*x̄*: Mean of gene x expression value in all cells

*w_i,j_*: Weight of a neighboring cell pair cell *i* and cell *j W*: Sum of all *w_i,j_*.

Squidpy [35] was used to calculate Moran’s I with the default settings.

## Results

### Overview of SC2Spa

SC2Spa is designed to infer spatial location of cells using deep learning (Fig 1A). SC2Spa is designed to map cells/spots to any type of ST data including image-based ST data [15], sequence based Visium [25], Slide-seqV2 [36] and Stereo-seq [37] data only based on the transcriptome. The core component of SC2Spa is a FCNN that stores the weights to map cells to their spatial location. For training, SC2Spa uses the transcriptome of ST data as the input and their spatial location of the ST data as the output. Once trained, SC2Spa accepts the transcriptome from ST and even independently generated scRNA-seq data to predict their spatial location (Fig 1A). Once the spatial coordinates are predicted, multiple cells or ST spatial indices can be mapped to a spatial index on the reference slide. For visualization, the multiple predictions within a certain range from the center of a spatial index are further merged with the weights considering the distance (Method).

**Figure 1.**
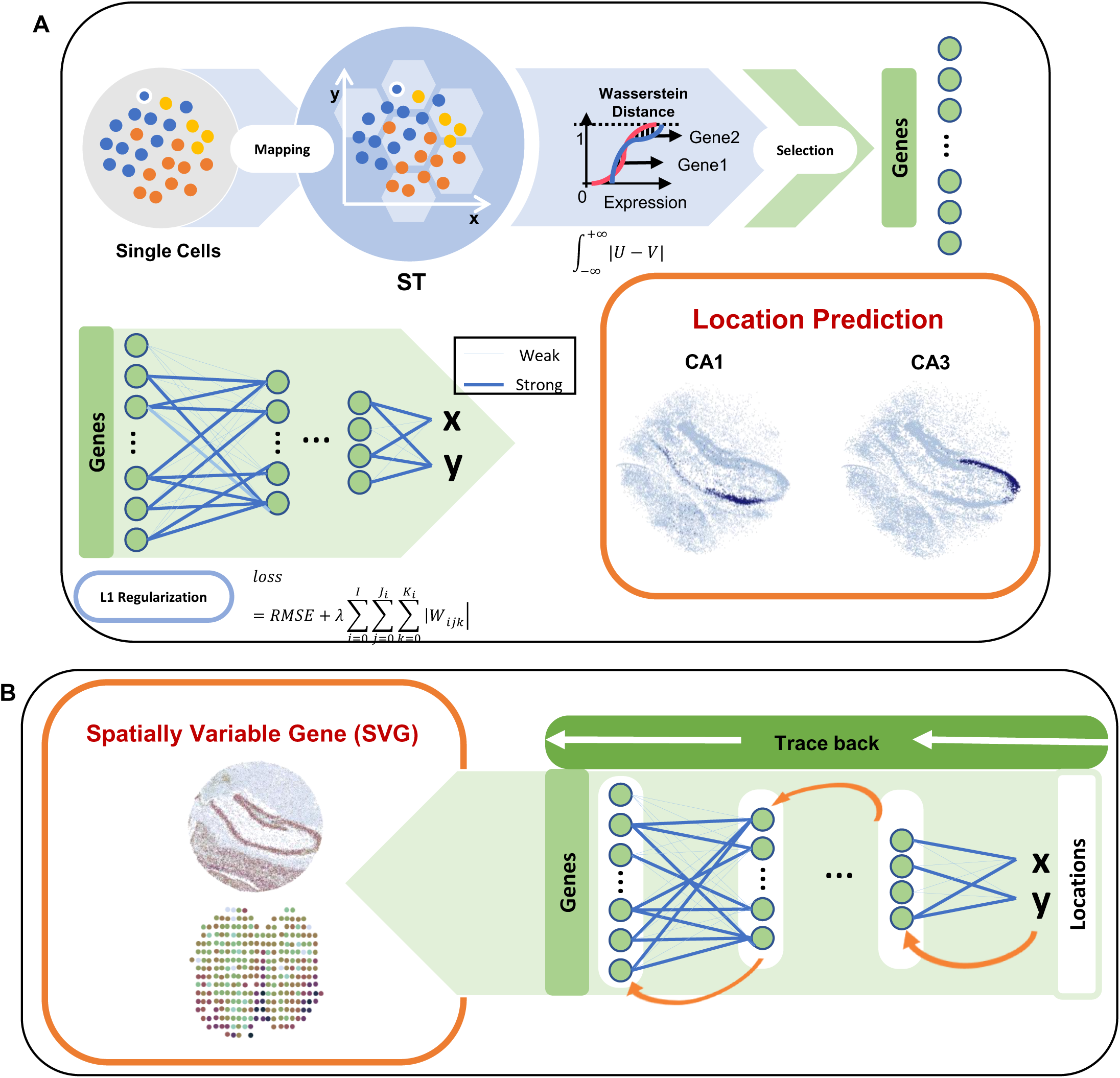
Overview of SC2Spa. **A.** SC2Spa is composed of a FCNN to map transcriptome to the spatial location. For training, SC2Spa uses the transcriptome from ST beads as the input and the spatial location of them on the ST assay as the output. The common genes within a specified range based on the Wasserstein distance are selected. The weights of the FCNN are constrained using L1 regularization to promoter sparsity. The trained FCNN is used to predict spatial location for the transcriptome from either ST beads or scRNA-seq data. The predicted location is further visualized on the ST assay. **B.** The weights of the FCNN are used to identify SVGs.

By default, SC2Spa uses 6 hidden layers and the number of nodes in these layers was set to decrease by 4 folds from 4096 (i.e. 4096, 1024, 256, 64, 16, and 4, respectively). A user can easily change the number of layers. The input layer is designed to accept the entire transcriptome of a cell or a spatial spot and the output layer is composed of 2 nodes for spatial coordinates. The rectified linear unit (ReLU) function is used for the first five hidden layers. To avoid overfitting, the input layer and the first 5 hidden layers were randomly deactivated during the training with a dropout rate of 0.05 and were normalized. The output layer uses the sigmoid function. The Adam algorithm [26], an algorithm for first-order gradient-based optimization of stochastic objective functions, was used to train the weights (Method). To provide a level of interpretability to the trained FCNN, SC2Spa trains the weights while penalizing the sum of weights by using L1 regularization (Method). The weights of the network that do not contribute to the location prediction tend to shrink to zero during training. The trained neural networks (NNs) provide additional interpretation about the ST data. SC2Spa determines the level of contribution for location prediction by backtracking the most informative weights (Fig 1B). The information is further used to determine spatially variable genes (SVGs).

### SC2Spa predicts the location in various ST datasets

First, we investigate the performance when testing with 10X Genomics Visium [25] data. Visium is a commercially available ST approach but with lower resolution (around 55μm in diameter per spot). For training we used brain data from a mouse (ID: ST8059049) and tested it using the Visium data from another mouse (ID: ST8059048). We visualized spatial mapping using the spot labels and the distance between the original and the mapped results (Fig 2A). While all methods revealed brain tissue architecture in the dataset (Fig 2A and Supplementary Figure 1), SC2Spa achieved the most accurate representation revealing regions such as Cornus Ammonis (CA) clearly While SpaOTsc showed a relatively good performance, mapping results on CA are sparse. The spots related with CA were scattered to other locations on that assay.

**Figure 2.**
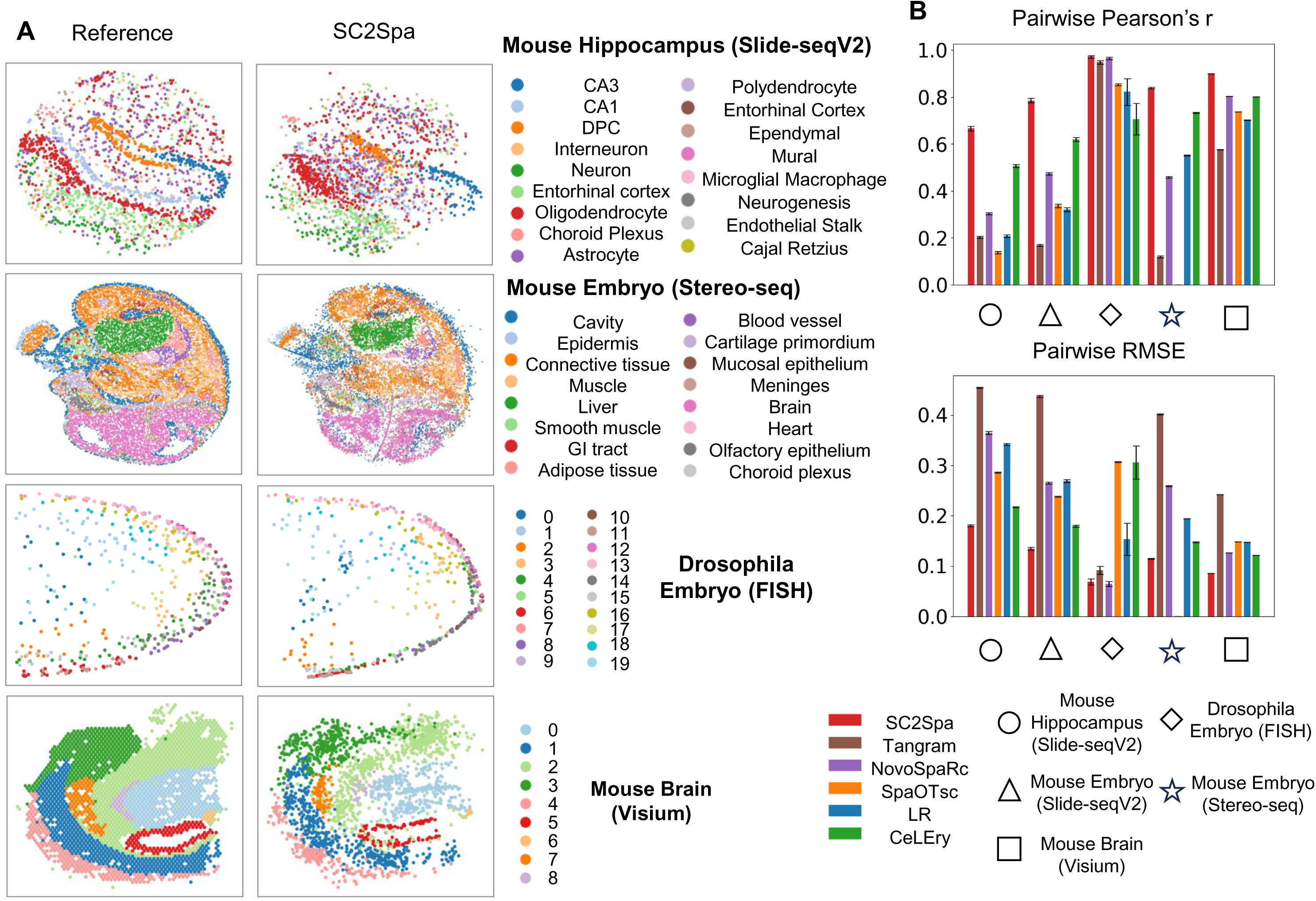
SC2Spa outperformed other predictors in predicting spatial origin from transcriptomics data when using independent data and cross-validation. **A**. The reference data and the predicted results from SC2Spa were visualized. The Visium data is trained and tested on the independent data from mouse brain. One of the five-fold cross-validation results is shown for the Slide-seqV2 mouse hippocampus data, Stereo-seq mouse embryo data and FISH drosophila embryo data. **B.** Comparison using pairwise Pearson’s *r* and pairwise RMSE. The bootstrap method was used to resample the spatial voxels 200 times to obtain error bars. For each bootstrap iteration, a sample of the same size as the original dataset was drawn with replacement.

To further assess the performance of SC2Spa in mapping spatial location, we employed a rigorous five-fold cross validation strategy. This involved testing it on diverse ST datasets including Slide-seqV2 from both the mouse brain and embryo [36], Stereo-seq data from mouse embryo [37], and in situ hybridization (ISH) data from drosophila embryos [15]. For benchmarking, we compared the performance against other location predictors including Tangram [18], Novosparc [16, 20], SpaOTsc [17], and CeLEry [23]. Additionally, we included a simple logistic regression (LR) [38].

For the Slide-seqV2 datasets in the mouse brain [36], which has a near cellular resolution, we included various cell types with the largest number of cells including tissue specific CA1 principal cells, CA2 principal cells, dentate principal cells and non-tissue specific interneurons, neurons, oligodendrocytes, and astrocytes (Fig 2A). After training, the coordinates for the transcriptome of the test set were visualized for all predictors (Fig 2A). The prediction by SC2Spa showed close similarity with the reference test set. SpaOTsc roughly recapitulated the CA composed of the CA1 and CA3 regions as well as the Dentate gyrus. However, many of the prediction by SpaOTsc were far from their original locations (Supplementary Figure 2). For example, SpaOTsc placed some predictions very close to the actual location of dentate principal cells (DPCs). However, it also generated other predictions significantly farther from the reference points (Supplementary Figure 3).

Other algorithms we tested failed in predicting the location of cells and could not retrieve the structure of the reference image (Supplementary Figure 2). CeLEry [23], another approach based on FCNN, predicted the rough location for CA3 and Dentate gyrus and other cell types as well. However, it could not recapitulate the general tissue patterns for these cells. Additional cross-validation tests using Slide-seqV2 from mouse embryo [36] further confirmed that SC2Spa and SpaOTsc accurately predicted the distribution of epithelial cells, myocytes and stromal cells while other approaches failed in predicting their distribution (Supplementary Figure 4).

Next, we performed 5-fold cross-validation using the Stereo-seq data from mouse embryo [37]. Stereo-seq is a sequencing-based ST data with a resolution (<1μm) higher than Slide-seq. We used the square-bin data (25μm). SC2Spa again recapitulated the reference location of the transcriptomes while other approaches failed in predicting them (Fig 2A). CeLEry captured the rough structure of the tissue architecture including the location of liver and brain. Tangram, NovoSpaRc showed far worse prediction performance (Supplementary Figure 5). Due to the extremely high memory requirements of SpaOTsc for large datasets, it was unable to run for this dataset.

To assess the performance using image based ST data, we used the three-dimensional (3D) fluorescence in situ hybridization (FISH) data from drosophila embryos [15]. These FISH data contains the transcriptomic profile of 84 genes from the drosophila embryo [15]. For this simulation, we prepared the neural network to predict 3D coordinates. In this 5-fold cross validation test, all approaches except for LR predicted the rough layout of fruit fly embryos (Supplementary Figure 6).

To enable quantitative assessment, we introduced two metrics: pairwise Pearson’s correlation coefficient (r) and pairwise root mean square error (RMSE). These metrics compare intercellular distances before and after mapping, independent of spatial rotation. By focusing on relative distances rather than absolute coordinates, they provide a robust and rotation-invariant evaluation of spatial relationship preservation (Method). For all the tests we performed, SC2Spa showed the least pairwise RMSE values (Fig 2B). The performance of CeLEry was next to SC2Spa in our tests. Even though the tissue structure appeared to be well identified by SpaOTsc in the image visualization, it did not get good scores in these metrics due to the frequent mapping to distal regions from the original location (Supplementary Figure 1-6).

### SC2Spa outperforms in mapping independent Slide-Seq data

Cross-validation was performed by splitting a single dataset into training and testing sets. We further evaluated SC2Spa’s performance using an independent dataset. We trained SC2Spa and other approaches using a Slide-seqV2 puck (ID: puck_200115_08) and tested using another puck (Query set, ID: puck_191204_01) from the mouse brain [36].

As an example, we show the beads for CA1 and CA3 pyramidal cells and the dentate gyrus for both the training and the query set (Fig 3). SC2Spa, trained on one puck, processed the transcriptome from the Query set, and predicted the location for the query data. The orientation of the predicted results followed the training set rather than the test set. Considering that the orientation from the query set is lost and only transcriptome is used as the input for SC2Spa, it is natural that the orientation of the predicted image follows that of the training set (Fig 3C).

**Figure 3.**
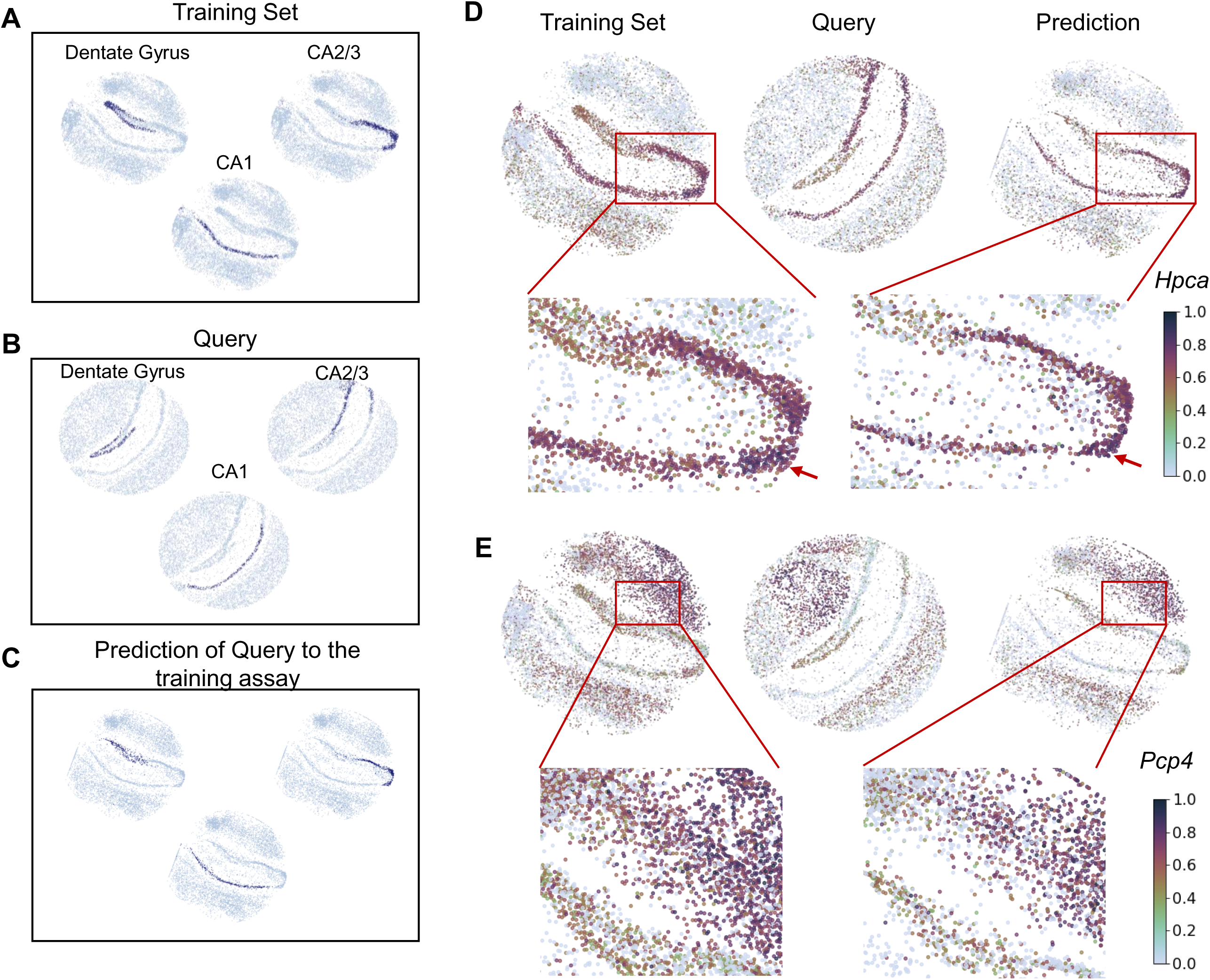
The trained SC2Spa recovers the locations for unseen ST data. **A.** The locations of dentate gyrus, CA1, CA2 and CA3 pyramidal cells in the training ST data (puck_200115_08) **B.** independent query set (puck_191204_01) and **C.** prediction. **D,E.** The spatial expression of *Hpca* and *Pcp4* in the training, and the testing sets as well as the predicted results. BVI is calculated for *Hpca* (0.35) and *Pcp4* (0.09) between the training set and the prediction. The color bar is to show the expression levels for the genes.

To assess the spatial similarity of the gene expression we used bivariate Moran’s I (BVI) [33, 34] (Method). BVI measures the spatial autocorrelation between two genes. If BVI=0, there is no spatial autocorrelation. Visualization of individual genes showed that the spatial distribution of genes such as *Hpca* (BVI = 0.35) and *Pcp4* (BVI=0.09) match well with the spatial patterns observed in the training as well as the Query set (Fig 3D and 3E). A deeper analysis revealed mild expression levels of *Hpca* in the Dentate Gyrus, which becomes stronger in CA regions with very strong expression levels in the CA2 region. These tendencies were observed in both the training set and the prediction images (Fig 3D). The gradual changes of the *Pcp4* expression were also observed in the training set and the predicted image (Fig 3E). The comparison also showed that SC2Spa excelled other approaches in location prediction (Supplementary Figure 7).

### SC2Spa maps single cell transcriptome to spatial locations at various resolutions

To further assess how spatial resolution impacts mapping accuracy, we evaluated SC2Spa’s performance using MERFISH data from a mouse brain (C57BL6J-1.080 as reference and C57BL6J-1.081 as query) [39]. We generated reference ST at two resolutions: 20μm and 50μm (Bin20 and Bin50, respectively) by merging the transcriptome of the reference cells. The cellular resolution query dataset was then used for testing. We found that SC2Spa maintained successful mapping of the MERFISH data regardless of the resolutions (Bin20 and Bin50) and outperformed other predictors. As the bin size increased (indicating lower resolution), the mapping performance mildly compromised (Supplementary Figure 8).

### SC2Spa predicts scRNA-seq spatial locations consistent with tissue anatomy

We further tested if SC2Spa can map scRNA-seq to Slide-Seq V2 and Visium data. First, we used SC2Spa trained with the cellular-resolution Slide-Seq V2 data from the mouse brain [36] and predicted the location of scRNA-seq using the publicly available data from the mouse brain [40]. The prediction results matched well with the known location for each cell type. For instance, the prediction for CA1, CA2, CA3 pyramidal cells and dentate gyrus granule cells matched well with the corresponding regions (Fig 4A). Genes such as *Nrgn* (BVI=0.28), *Fibcd1* (BVI=0.27), *Tshz2* (BVI=0.29), and *Hs3st4* (BVI=0.26) confirmed the similarity between the reference ST data and the prediction (Fig 4B).

**Figure 4.**
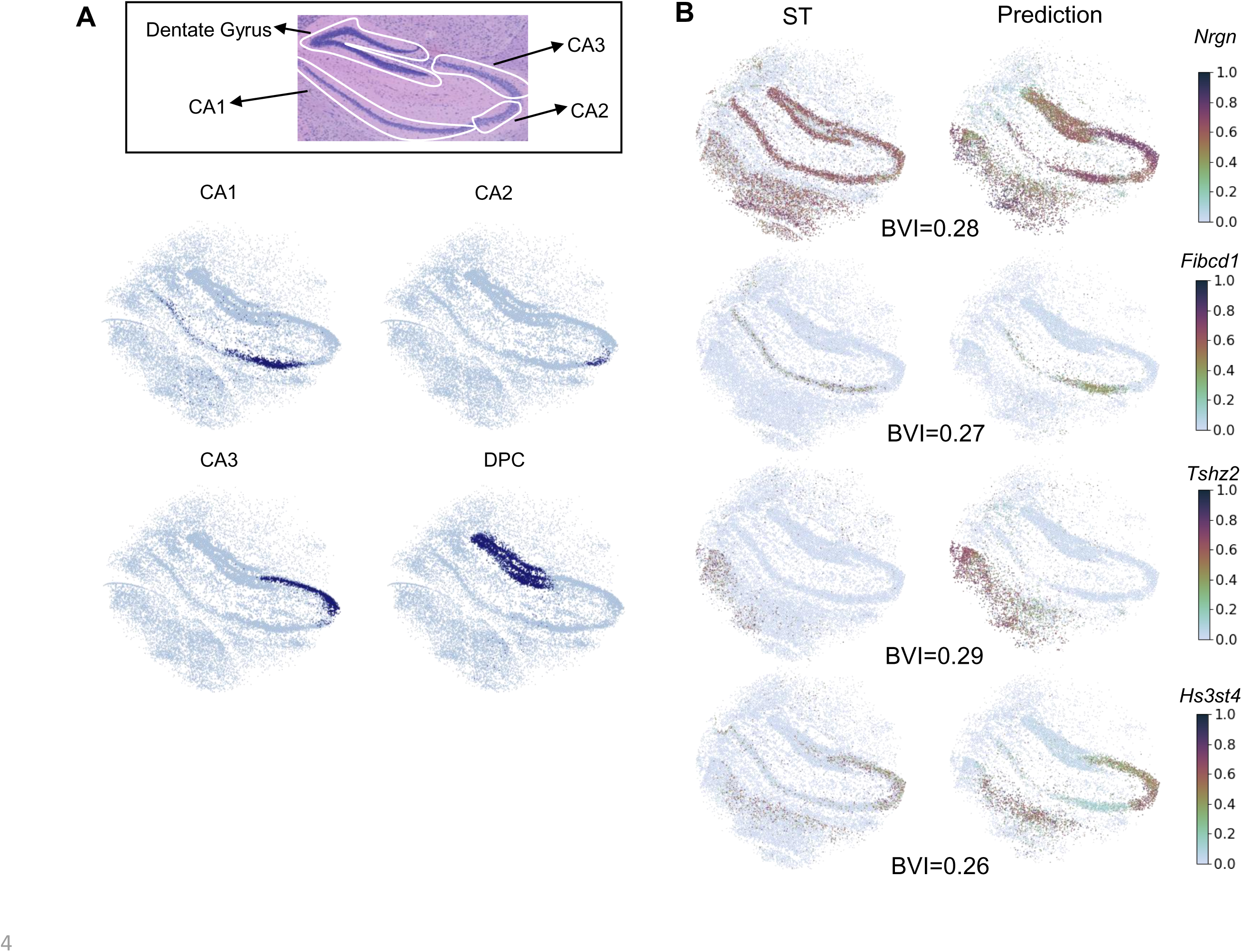
SC2Spa predicted the locations for scRNA-seq data. **A.** The predicted locations for CA1, CA2, CA3 pyramidal cells and dentate principal cells from scRNA-seq. The histology image is from [27] and is manually labeled. **B.** Spatial expression patterns of *Nrgn*, *Fibcd1*, *Tshz2* and *Hs3st4* show the similarity between the reference and the predicted results. BVIs were calculated for these genes between the reference ST and the prediction by SC2Spa.

We further evaluated if SC2Spa can still be used to predict the location of single cells even when trained on low-resolution Visium data. We used public Visium 10X for training and single nucleus RNA sequencing (snRNA-seq) datasets obtained from 2 mouse brains for testing (one from the same mouse and another from the other mouse) [27]. We assessed the location mapping of oligodendrocytes (localized mainly in the white matter (WM)), inhibitory neurons (mainly in hypothalamus (HYP)), and excitatory neurons (mainly in thalamus (THA)) that express marker genes such as *Mbp*, *Gad2*, and *Slc17a6,* respectively [27] (Fig 5A). The location mapping of the scRNA-seq from the same mouse brain exhibited the cell localization patterns matching with our prior knowledge (Fig 5B). The test using the scRNA-seq from an independent mouse brain also showed similar patterns (Fig 5C). Genes such as *Rgs9* (BVI=0.2), *Eps8l2* (BVI=0.16), *Stard8* (BVI=0.05), and *Otof* (BVI=0.14) further confirmed the similarity between the predicted results and the Visium data (Fig 5D).

**Figure 5.**
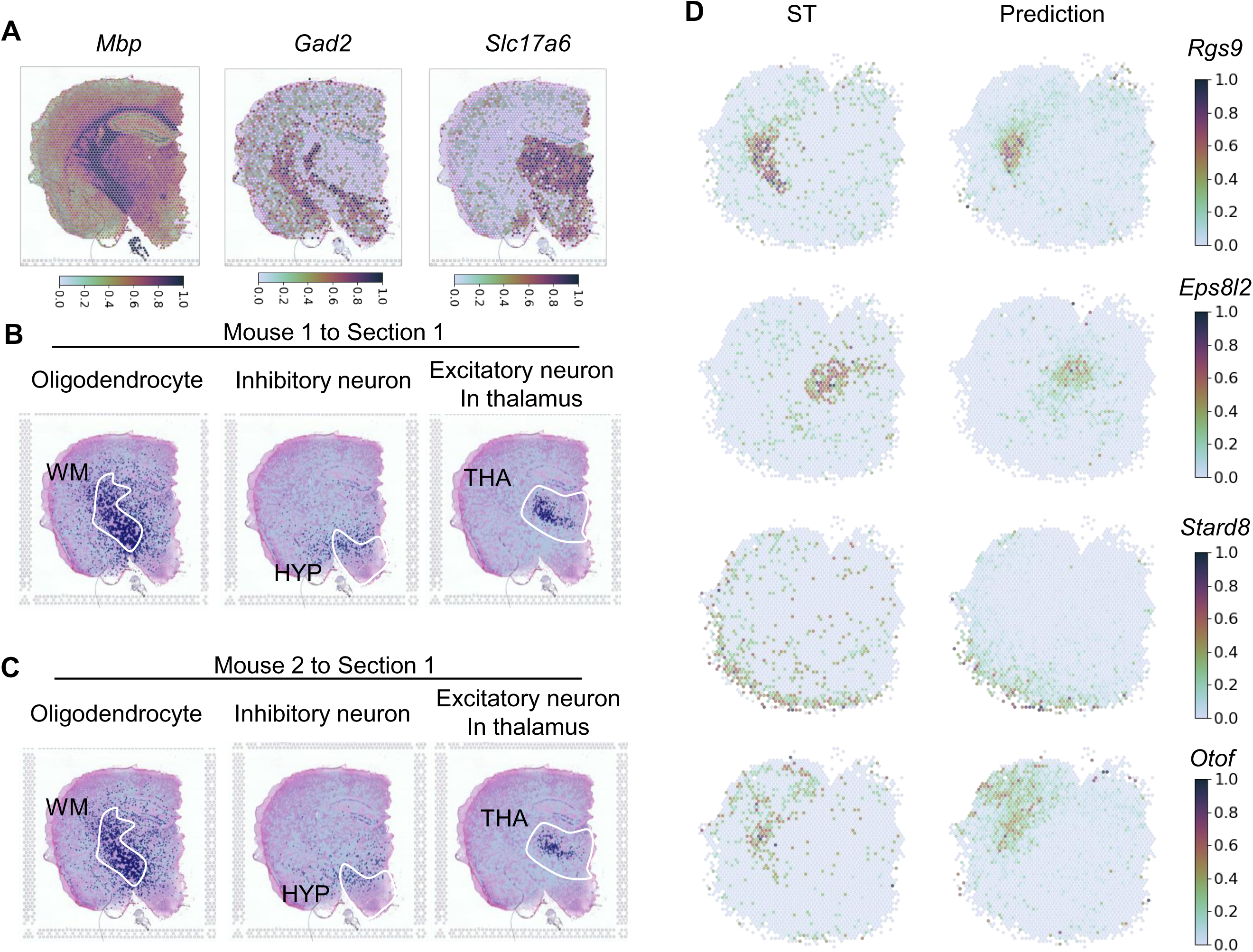
SC2Spa trained on Visium data predicted the locations for scRNA-seq. **A.** The spatial expression profiles of *Mbp*, *Gad2*, and *Slc17a6* on the Visium data. **B.** The predicted localization of oligodendrocytes, inhibitory neurons and excitatory neurons in thalamus from the scRNA-seq from the same mouse brain. **C**. Spatial reconstruction using the scRNA-seq data from an independent mouse brain. **D.** The expression of *Rgs9* (BVI=0.2), *Eps8l2* (BVI=0.16), *Stard8* (BVI=0.25), and *Otof* (BVI=0.14) in the reference data and the predicted results.

We further applied SC2Spa to the human kidney cancer Visium data and mapped the location using the accompanying scRNA-seq data [28]. SC2Spa successfully captured the spatial gene expression patterns in the kidney cancer tissue. For instance, *DMKN* (BVI = 0.32), *EGR1* (BVI = 0.18), *KCNJ1* (BVI = 0.35), *MT1H* (BVI = 0.24), *RASD1* (BVI = 0.23) and *RPL37* (BVI = 0.20) show high similarities in spatial expression between the reference Visium data and prediction (Supplementary Figure 9). We also trained SC2Spa using human breast cancer Visum data [29] and tested the mapping performance using the two scRNA-seq datasets derived from formalin-fixed paraffin-embedded samples (scFFPE-seq). These datasets were generated using the 3’ and 5’ protocols provided by 10X Genomics, respectively. SC2Spa demonstrated consistent performance regardless of the scRNA-seq protocol employed. This is exemplified by genes like *CYB561, FASN*, and *PVALB*, which exhibited high spatial similarity between the reference and both predicted locations (Supplementary Figure 10).

### SC2Spa predicts SVGs

SC2Spa has the ability to score the SVGs from their weight space. We compared the SVGs obtained with SC2Spa to those obtained by SpatialDE [41]. SpatialDE uses statistical tests to identify genes with spatial variable expression patterns and calculates the log-likelihood ratio (LLR) score against a random distribution. We plotted the SpatialDE’s LLR values against the scores indicating contribution to location prediction from SC2Spa and found positive relationships (Supplementary Figure 11A). Genes such as *Ttr*, *Pcp4*, *Calb2* and *Tshz2* showed a high score for both of the methods (Supplementary Figure 11B). We also found genes such as *Gm13872*, *Gm26724,* and *2010005H15Rik* were only observed by SpatialDE (Supplementary Figure 11C). Moran’s I has been used to assess the distribution of spatial gene expression [35, 42]. We found that the scores for contribution to location prediction by SC2Spa better correlate with Moran’s I than SpatialDE scores (Supplementary Figure 11D). Visual inspection showed that SpatialDE calculated high scores for rarely expressed genes even though their absolute Moran’s I score was less than 0.001(Supplementary Figure 11C). Similar outcomes were observed in the second Slide-seqV2 mouse hippocampus dataset and a Slide-seqV2 mouse embryo dataset (Supplementary Figure 12) and (Supplementary Figure 13). Some genes (*Tert*, *Gm19684*, and *Gm10640* for mouse hippocampus and *Krt75*, *Tmem171*, and *Has1* for mouse embryo) with a high SpatialDE score did not exhibit any spatial pattern and had an absolute Moran’s I score of less than 0.0001. Our results show that SC2Spa can suggest an alternative way to obtain SVGs.

## Discussion

By providing both transcriptomic profiles and spatial location, ST is becoming more popular for many areas in biology [43–47]. However, there are still some limitations in using ST data. Imaging-based ST techniques such as seqFISH+ [48], MERFISH [49] and EEL FISH [50] can obtain cellular resolution ST data. However, they are limited by the number of genes they capture simultaneously. Sequencing-based STtechniques target RNA species unbiasedly. However, a spatial barcode can be mapped to multiple cells or a portion of a cell depending on the resolution [25, 36, 37, 51–55]. Even without the spatial component, scRNA-seq is still useful as it provides an unbiased transcriptome measure of cell. There are also sequencing-based ST approaches with even sub-cellular resolution. The quality of these high-resolution spatial transcriptomics data such as Visium HD and Stereo-seq [37] are still not enough so that they do not provide the quality near to scRNA-seq data. Therefore, they often generate a near Visium resolution dataset for subsequent analysis [56]. Even after cell segmentation followed by profiling transcriptome per cell, the quality is still poorer than scRNA-seq. SC2Spa can be used to refine and enhance the data obtained from these newer methods, potentially bridging the gap between spatial resolution and data quality. Cellular resolution approaches such as Xenium are still limited in the number of genes to use. scRNA-seq mapping can be used to impute missing genes. Mapping by SC2Spa can impute missing genes. Plenty of resources have been accumulated in the form of scRNA-seq. Location prediction for scRNA-seq will be useful to explain cell heterogeneity, communication and function in a given tissue architecture. SC2spa is best suited for applications where the query dataset similar to the reference tissue. SC2Spa cannot perform well on tissues that contain cell types it was not trained on. SC2Spa has been tested on various datasets. However, in theory, it may not perform well with image-based spatial transcriptomics technologies that rely on a limited number of cell type marker genes, as these datasets lack diverse gene expression information related to spatial positioning.

SC2Spa not only excels in cross-validation tests, but it also demonstrates performance in predicting the spatial location of scRNA-seq data, even when trained on low-resolution Visum data. Experiments utilizing binned MERFISH data demonstrate SC2Spa’s ability to map transcriptomes even to lower resolution data, albeit with a gradual decline in performance as the resolution coarsens (Supplementary Figure 8). ST data such as Visium have the compositional profiles coming from multiple cell types. Our test show that SC2Spa performs well even when it was trained on the compositional data. SC2Spa may have captured genes associated with global structure, cell type specific genes and cell-cell communication related genes remaining in the compositional data. There are a number of methods to decompose spatial beads into the cell type proportions composing the transcriptomic profile [57–63]. Our approach is different in that we try to find the best position that a cell can be mapped based on transcriptome information and do not aim to decompose spatial transcriptome.

Deep learning often suffers from overfitting. To evaluate the potential for overfitting in SC2Spa, we employed 5-fold cross-validation using Slide-seqV2 mouse hippocampus data. During SC2Spa training, we monitored RMSE for both the training and test sets (Supplementary Figure 14). As expected, the training RMSE steadily decreased, indicating the model’s ability to learn the training data. A general downward trend in the test RMSE signifies that overfitting does not severely compromise SC2Spa’s performance.

CeLEry [23], another deep learning-based approach, showed a comparable performance to SC2Spa especially for Visium data (Supplementary Figure 1, Supplementary Figure 2, Supplementary Figure 4, Supplementary Figure 5, and Supplementary Figure 6), indicating the usefulness of deep-learning for this tasks. Despite successful mapping of gene expression, the predicted locations lacked precision, making it difficult to clearly define tissue boundaries. Interestingly, CeLEry underperformed on 3D FISH data, which is considered easier for spatial mapping (Supplementary Figure 5). We further investigated the factors that influence the performance of SC2Spa over CeLEry. CeLEry used L2 regularization (weight: 1e-5) with 3 hidden layers for NN while SC2Spa used 6 hidden layers with L1 regularization by default. We found that the performance becomes enhanced as the number of layers increased (Supplementary Figure 15). We also found that L1 produced a better performance. The combination of these factors may influence the performance of SC2Spa. Besides, CeLEry uses data augmentation, which may not be useful for spatial mapping particularly for sequencing-based data that already has a sufficient amount of data.

We add a penalty on weight magnitudes to promote sparsity in learned features. Unlike dropout, which prevents overfitting by randomly deactivating neurons, our penalty encourages zero weights for uninformative genes, improving interpretability. This aids feature selection, reduces dimensionality, and filters noisy gene expression more effectively than dropout.

We introduce a penalty term on the absolute values of weights to promote sparsity in the model’s learned representations. This penalty encourages the model to focus on a smaller subset of informative genes, reducing noise and enhancing interpretability. While a standard neural network with dropout provides regularization by randomly deactivating neurons during training, it does not explicitly enforce sparse weight distributions. In contrast, our sparsity-inducing penalty drives the model to assign low or zero weights to less relevant features, reducing dimensionality in a biologically meaningful way. This approach enhances feature selection and interpretability by identifying genes most relevant to spatial mapping.

The trained weights also provide additional knowledge about the genes. SC2Spa scores the genes according to the contribution towards location prediction. We regarded them as SVGs even though the weights were not originally designed to. The SVGs from SC2Spa better correlated with Moran’s I than those recovered by SpatialDE [41].

SC2Spa is designed to accept one ST data. It is possible to use multiple ST datasets if multiple ST data are merged when the ST data have similar spatial expression patterns and share a similar coordinate system. In our tests, SC2Spa, Tangram and logistic regression were run with a 24GB RTX Titan GPU. The other CPU-based softwares were run with an 18-core Intel® Xeon(R) W-2195 CPU of 2.30GHz. We compared SC2Spa with existing algorithms for the applications where the locations of the query data are available (Supplementary Table 1). As SC2Spa maps gene expression directly to coordinates the computational complexity of SC2Spa increases linearly to the size of reference data. The complexity of optimal transport-based methods increases linearly with the product of the magnitudes of the reference and query data. When both the quantity of the reference data and the query data increases, the complexity of the other methods increases quadratically. As the size of the input data increases, SC2Spa will exhibits a distinct advantage over other algorithms in terms of execution time (Supplementary Figure 16, Supplementary Table 2).

The mapping of scRNA-seq into Visium can suggest potential use of SC2Spa in enhancing the spatial resolution of Visium data. scRNA-seq has been used to study cell-cell communication [64, 65]. However, it is of note that additional spatial information can boost the performance of predicting cell-cell communication [66]. We applied SpaTalk [66], a software to detect cell-cell communication from ST data, using the spatial information provided by SC2Spa. We found the interaction of interneuron and astrocyte through ErbB4 and Hbegf in the spatial domain (Supplementary Figure. 17).

### Conclusions

We developed a new framework SC2Spa to map scRNA-seq to spatial locations. Both high-resolution and low-resolution spatial transcriptomics (ST) references can be used for the spatial mapping with SC2Spa.The location prediction by SC2Spa demonstrates the emergence of expression patterns contributing tissue architecture contained within scRNA-seq. SC2Spa also provides functions for the identification of spatially variable genes. SC2Spa enhances the resolution of low-resolution spatial transcriptomics data, imputes missing values, improves spatial gene expression reconstruction, infers spatial organization from scRNA-seq when ST data is unavailable, and provides deeper insights into cell-cell communication.

## List of abbreviations

ST: Spatial Transcriptomics
ScRNA-seq: single cell RNA sequencing
FCNN: fully connected neural network
SVG: spatially variable genes
UMI: unique molecular identifier
CDF: cumulative distribution function
RAM: random access memory
ReLU: rectified linear unit
NNs: Neural networks
CA: Cornus Ammonis
ISH: In situ hybridization
LR: Losgistic regression
DPCs: Dentate principal cells
FISH: Fluorescence in situ hybridization
RMSE: Root mean square error
BVI: Bivariate Moran’s I
SnRNA-seq: Single nucleus RNA sequencing
WM: White matter
HYP: Hypothalamus
THA: Thalamus
ScFFPE-seq: Single cell formalin-fixed paraffin-embedded sequencing
LLR: Log-likelihood ratio

## Declarations

### Ethics approval and consent to participate

Not applicable

### Consent for publication

Not applicable

### Availability of data and materials

SC2Spa is available at https://github.com/linbuliao/SC2Spa. The analysis for this manuscript is available at https://github.com/linbuliao/SC2Spa_Notebooks. The data and trained model for running SC2Spa is deposited at Figshare: https://doi.org/10.6084/m9.figshare.21829905. All data and analysis involved in this manuscript is deposited at Zenodo: https://zenodo.org/records/8252715. We also provide well-structured tutorials on our ReadTheDocs website: https://sc2spa.readthedocs.io/en/latest/.

The data used in this manuscript were obtained from the following sources: Slide-seqV2 mouse embryo and hippocampus data: https://singlecell.broadinstitute.org/single_cell/study/SCP815/sensitive-spatial-genome-wide-expression-profiling-at-cellular-resolution#study-summary

ScRNA-seq mouse hippocampus data (applied for mapping to the Slide-seqV2 hippocampus dataset): http://dropviz.org/

Stereo-seq mouse embryo data: https://db.cngb.org/stomics/mosta/download/

Visium mouse brain data (Cell2location): https://www.ebi.ac.uk/biostudies/arrayexpress/studies/E-MTAB-11114

SnRNA-seq mouse brain data (Cell2location): https://www.ebi.ac.uk/biostudies/arrayexpress/studies/E-MTAB-11115

3D FISH drosophila embryo: https://shiny.mdc-berlin.de/DVEX/

MERFISH mouse brain data: https://cellxgene.cziscience.com/collections/0cca8620-8dee-45d0-aef5-23f032a5cf09

Visium human kidney cancer and paired snRNA-seq data: https://data.mendeley.com/datasets/g67bkbnhhg/1

Visium human breast cancer and paired scFFPE-seq data: https://www.10xgenomics.com/products/xenium-in-situ/preview-dataset-human-breast

### Authors’ contributions

K.J.W, R.G. and L.L. conceived the project. L.L. developed the software. L.L. performed data analysis. H.K., K.J.W., G.K., J.K.and C.H.B. helped with data analysis. E.M., AMP, AK, PB, RW, JT, PF, RG carried out experiments. R.G. and E.M. helped with the experimental details. All wrote the manuscript.

## Acknowledgements

Not applicable

