## Supplementary Figures for "SC2Spa: a deep learning based approach to map transcriptome to spatial origins at cellular resolution"

Supplementary File

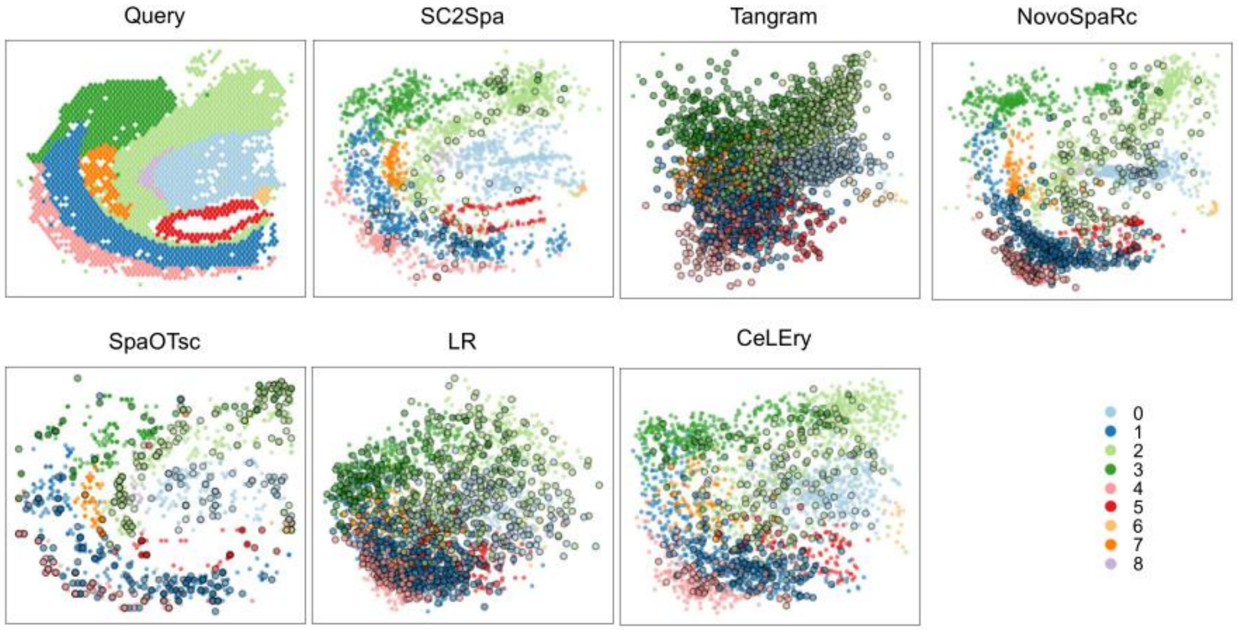


**Supplementary Figure 1. Performance comparison when testing using an independent Visium mouse brain dataset.** The Visium data used for Query and the predicted results. From the Query we obtained the clustering information. The beads for 8 clusters were visualized. When mapping two spatial sections, we calculated the average pairwise Euclidean distance difference between predicted and true values for each spot in the query section. Cells with a significantly large average pairwise Euclidean distance were marked with black circles.

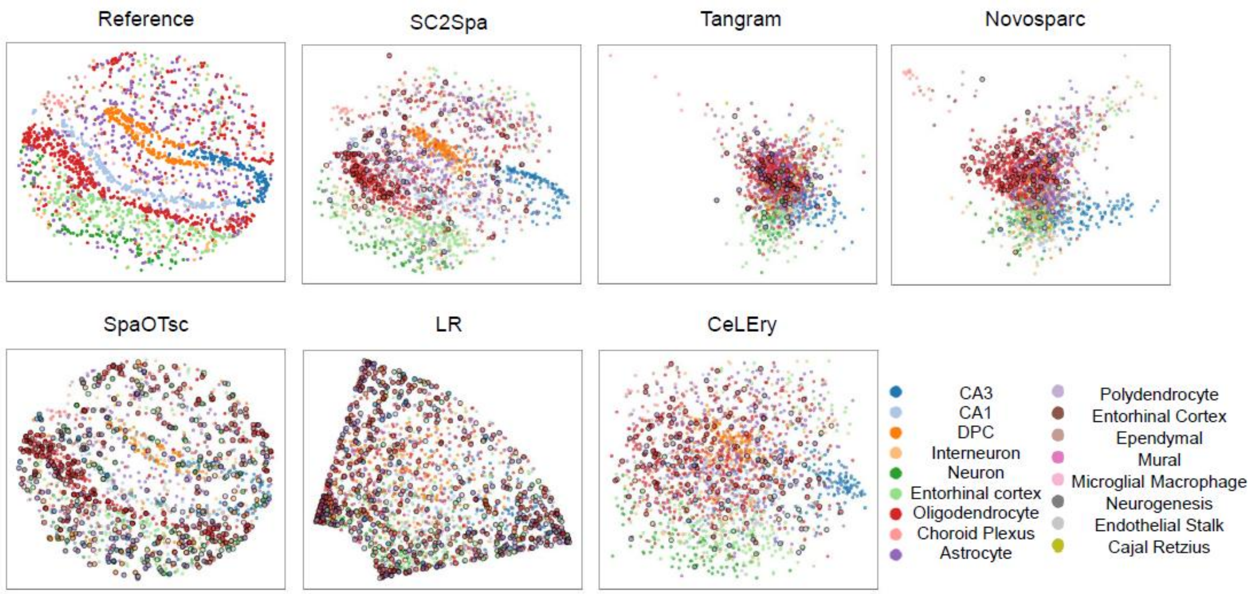


**Supplementary Figure 2. The reference image data and the predicted location using a SlideSeq-V2 mouse brain.** The reference image and the predicted location for SC2Spa, Tangram, Novosparc, SpaOTsc, LR and CeLEry were visualized for the leave-out beads for various cell types including CA1/3 principal cell, dentate principal cell, Interneuron, neuron, entorhinal cortex, oligodendrocyte and astrocyte. Cells that are beyond a certain distance from their original locations were outlined with a black circle and drawn larger. The cut-off distance was set at the 96th percentile of the Euclidean distance between predicted by SC2Spa and true spatial coordinates.

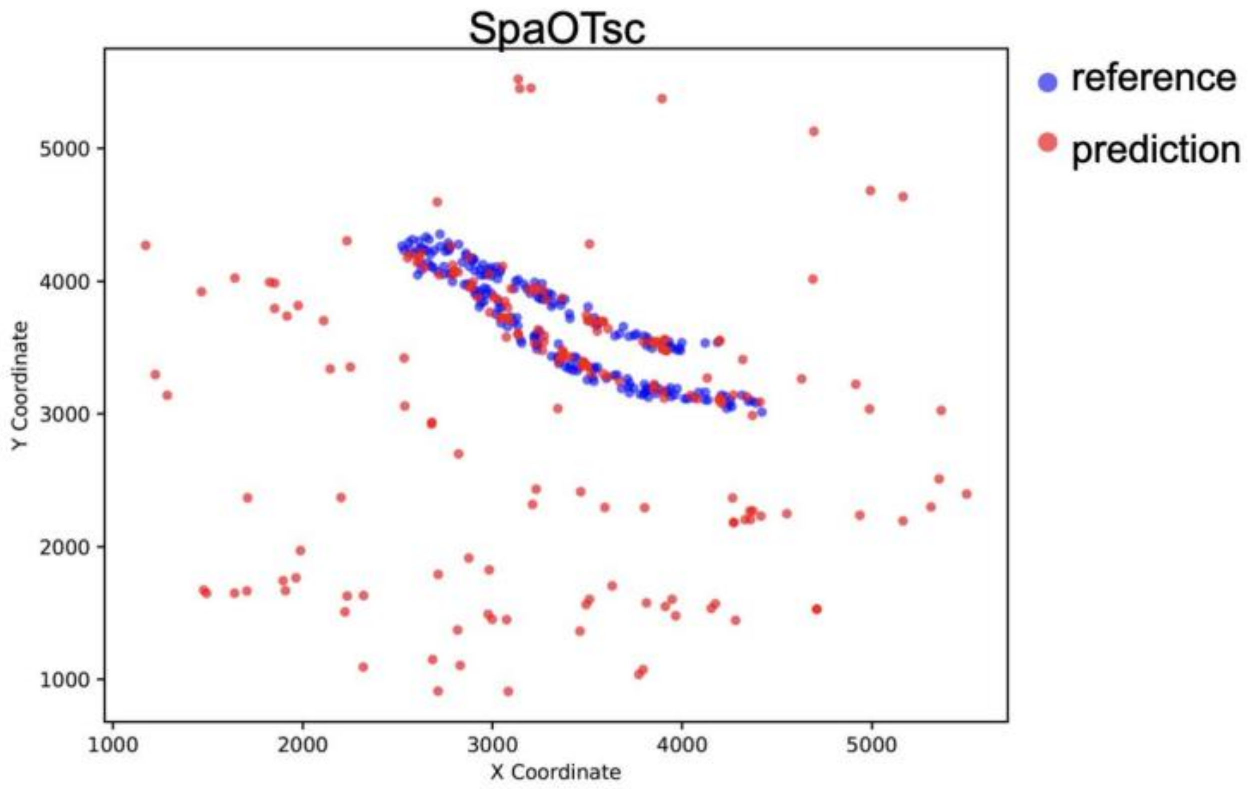


**Supplementary Figure 3. Location prediction of DPCs by SpaOTsc**. We visualized the prediction of SpaOTsc for DPCs.

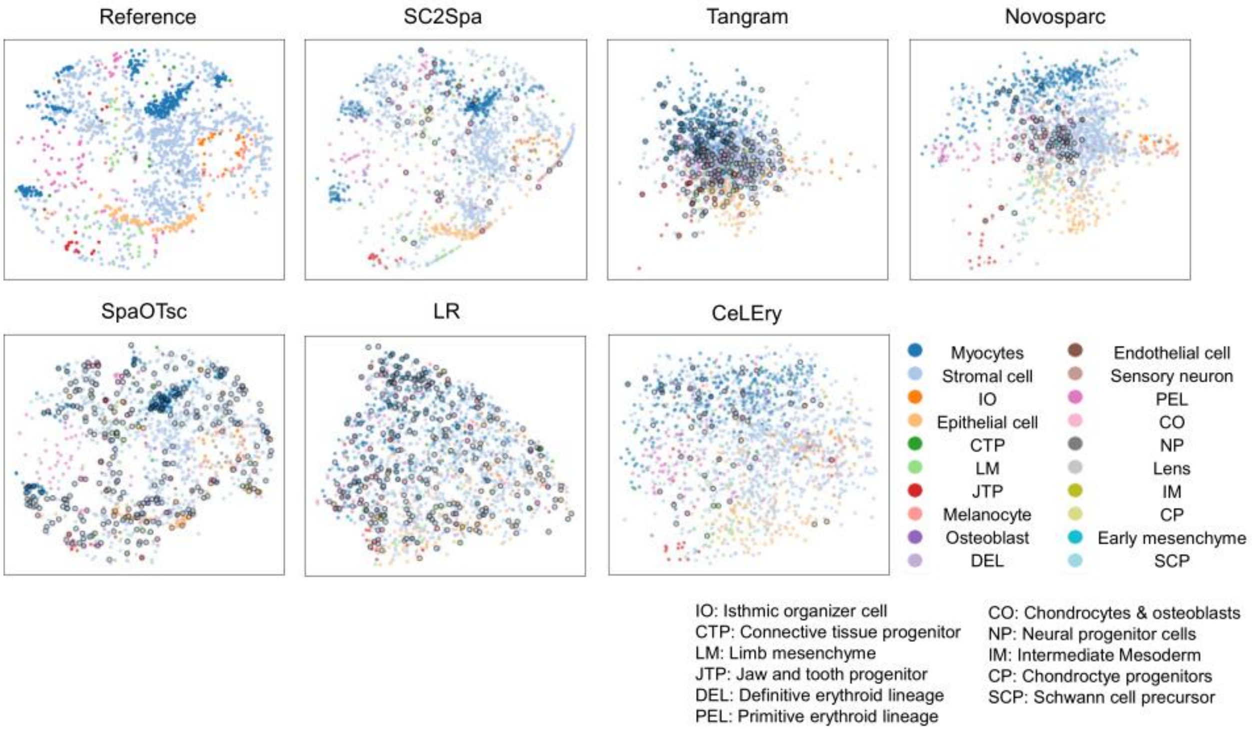


**Supplementary Figure 4. The reference image data and the predicted location using a SlideSeq-V2 mouse embryo.** The reference image and the predicted location for SC2Spa, Tangram, Novosparc, SpaOTsc, LR and CeLEry were visualized for the leave-out beads for 20 cell types including myocytes, stromal cells, isthmic organizer cells, epithelial cells, limb mesenchymal call, jaw and tooth progenitors, melanocytes and primitive erythroid lineage cells. Cells far from their original locations by a certain distance were highlighted with a black circle. The cut-off distance was set at the 96th percentile of the Euclidean distance between predicted by SC2Spa and true spatial coordinates.

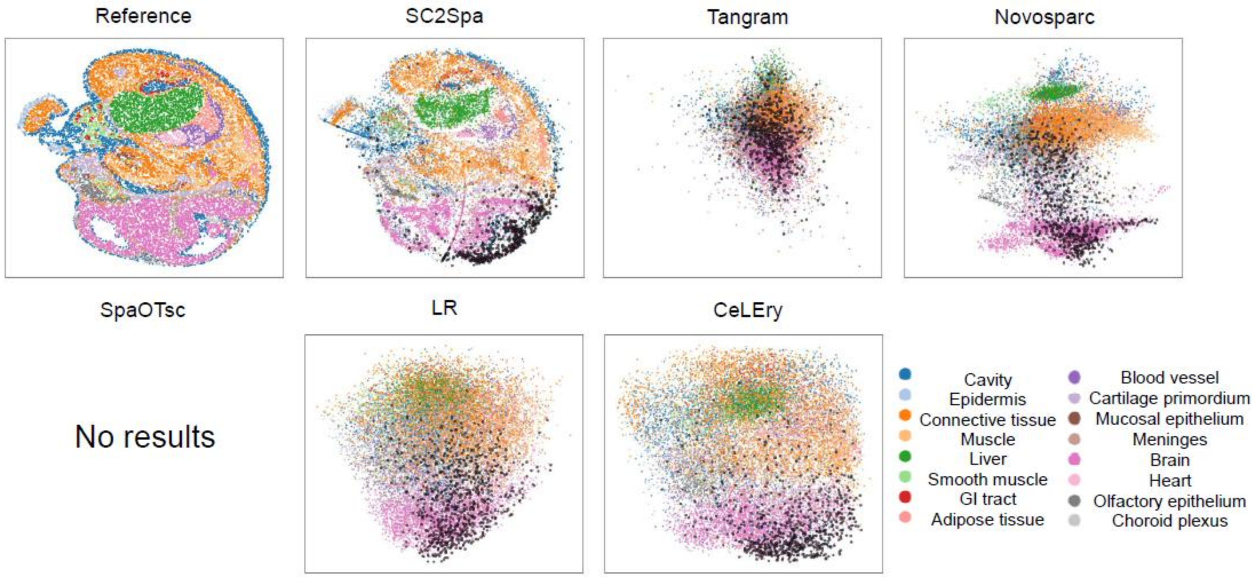


**Supplementary Figure 5. The reference image data and the predicted location of SC2Spa and other algorithms for a Stereo-seq datasets.** The reference image and the predicted location for SC2Spa, Tangram, Novosparc, LR and CeLEry were visualized for **A**. square-bin data from mouse embryo and **B**. cell-bin data from mouse brain (E16.5). Predictions that were far from a certain distance from their original locations were highlighted with a black circle. The cut-off distance was set at the 96th percentile of the Euclidean distance between predicted by SC2Spa and true spatial coordinates. SpaOTsc cannot run on this large size dataset due to its memory requirement.

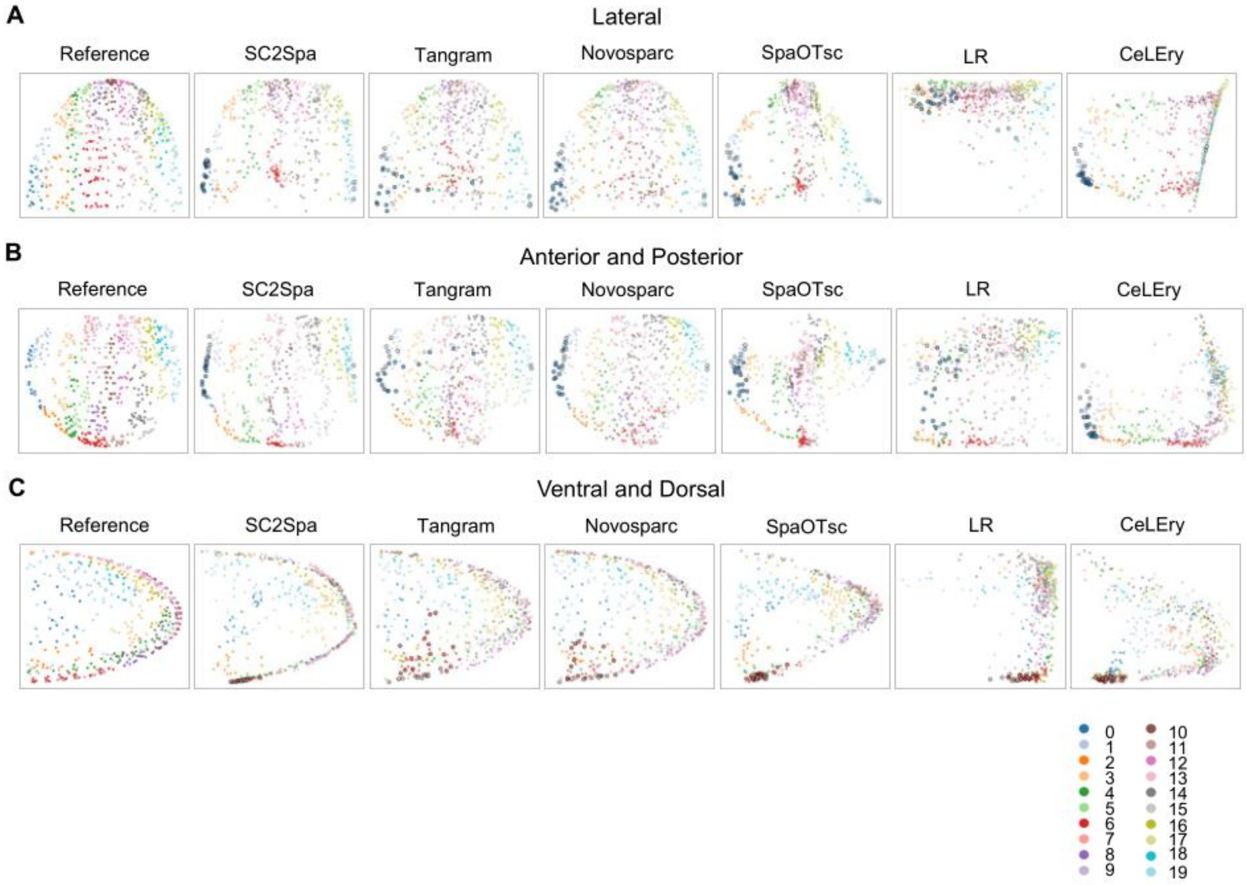


**Supplementary Figure 6. The reference and predicted locations using the 3D FISH drosophila embryo data.**

The original and predicted locations of ST beads in eight clusters were represented graphically. The drosophila embryo beads retained three-dimensional space information. The beads were viewed from three angles: lateral, anterior and posterior, as well as ventral and dorsal. Predictions that were far from a certain distance from their original locations were highlighted with a black circle. The cut-off distance was set at the 96th percentile of the Euclidean distance between predicted by SC2Spa and true spatial coordinates.

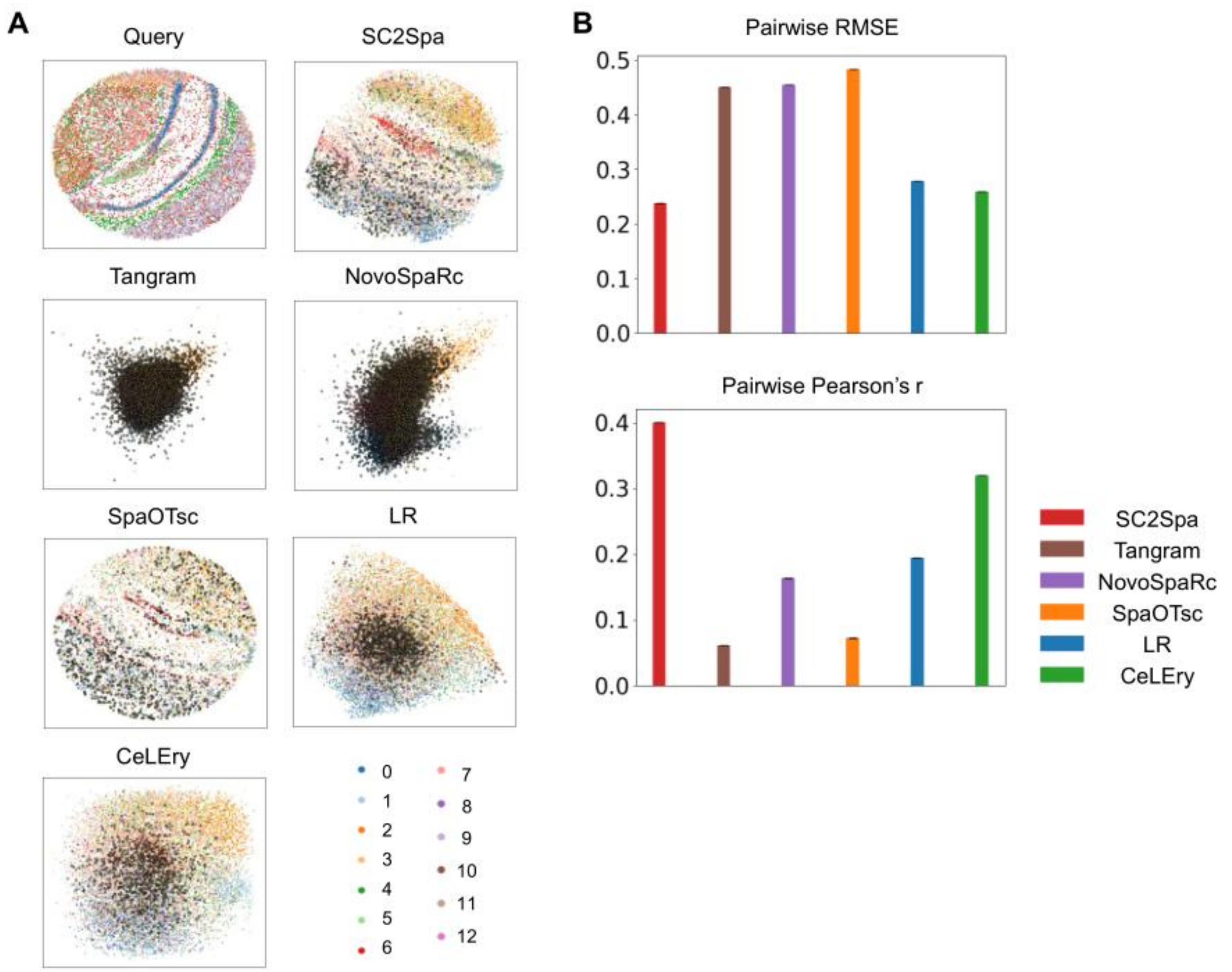


**Supplementary Figure 7. Performance comparison when testing with an independent Slide-seqV2 mouse brain dataset. A.** The Query SlideSeqV2 data and the predicted results. **B.** Comparison of the pairwise RMSE and the pairwise Pearson's *r*. When mapping two spatial sections, we computed the average difference in pairwise Euclidean distances between predicted and true values for each spot in the query section. Predictions that were far from a certain distance from their original locations were highlighted with a black circle. The cut-off distance was set at the 96th percentile of the Euclidean distance between predicted by SC2Spa and true spatial coordinates.

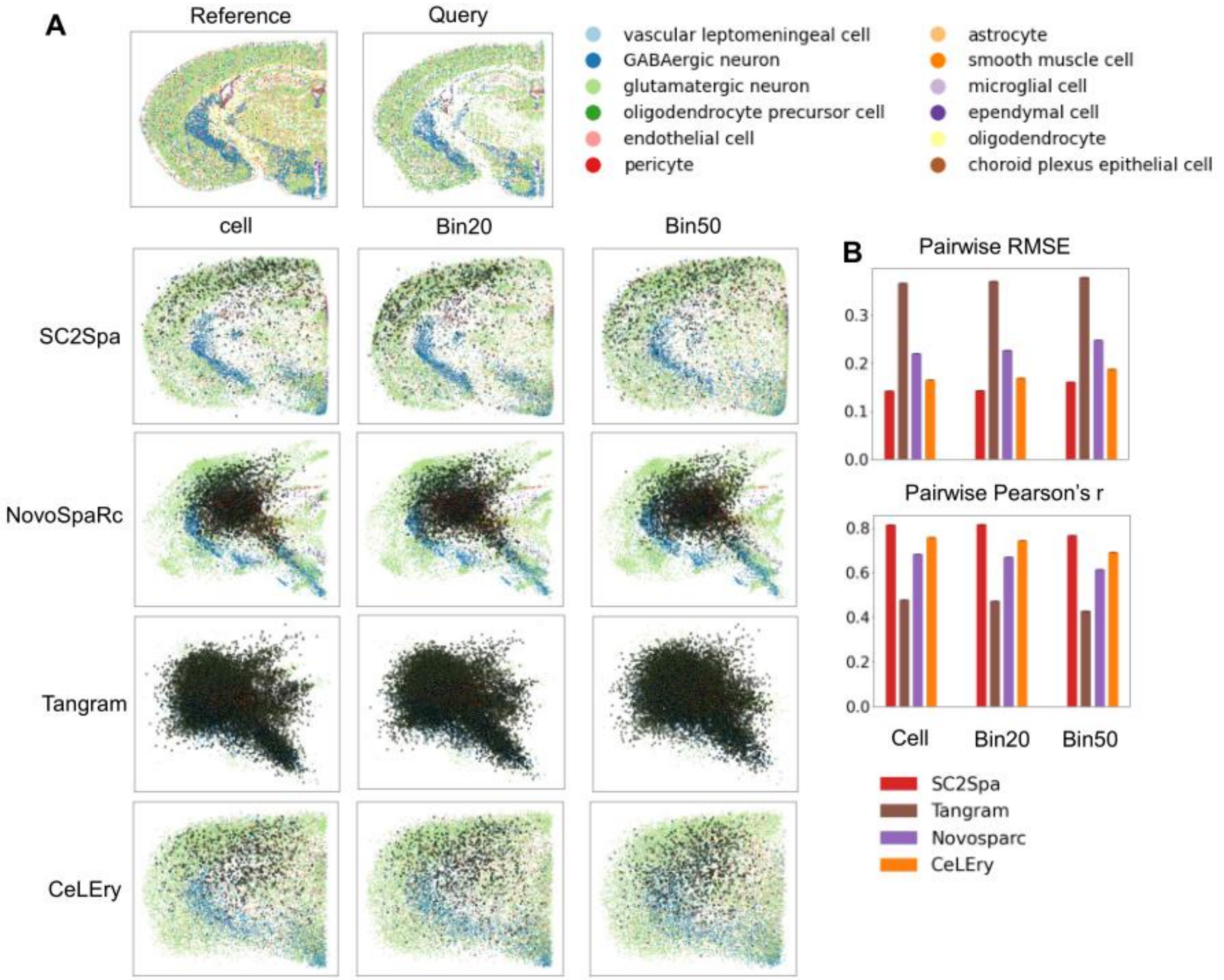


**Supplementary Figure 8. Performance comparison for mapping MERFISH data onto various resolution. A.** mapping results onto used cellular, 20μm bin (Bin20) and 50μm bin (Bin data) data. **B.** Comparison of the pairwise RMSE and the pairwise Pearson's *r*. During the mapping of two spatial sections, we calculated the average pairwise Euclidean distance difference between predicted and true values for each spot in the query section. Predictions that were far from a certain distance from their original locations were highlighted with a black circle. The cut-off distance was set at the 96th percentile of the Euclidean distance between predicted by SC2Spa and true spatial coordinates. SpaOTsc cannot run on this dataset.

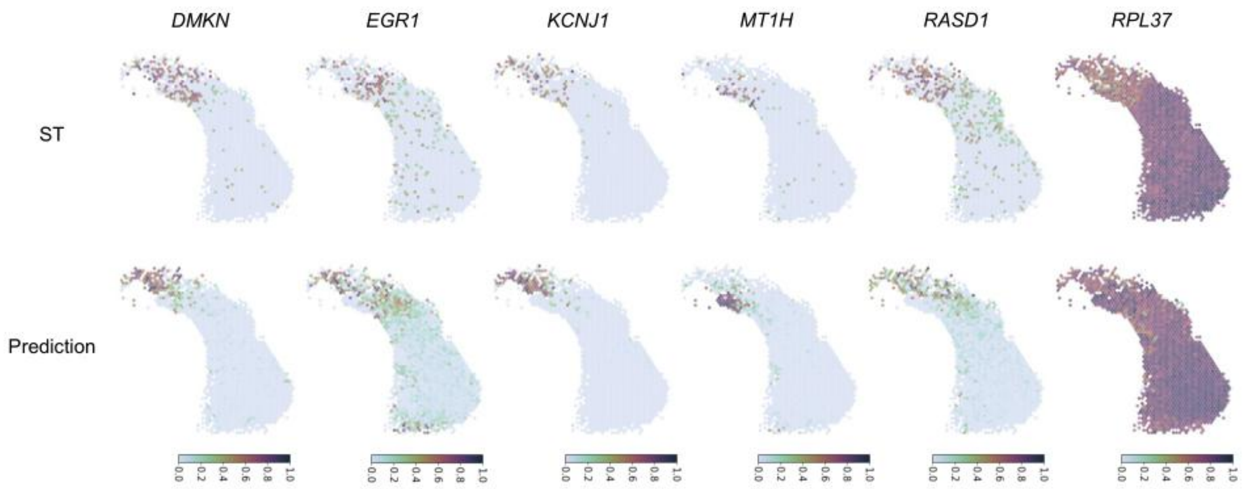


**Supplementary Figure 9. SC2Spa predicts scRNA-seq spatial locations with high consistency to reference Visium data.** SC2Spa is trained using the Visium dataset for human kidney cancer. The spatial expression for *DMKN, EGR1, KCNJ1, MT1H, RASD1, RPL37* is visualized in the reference ST data and the prediction.

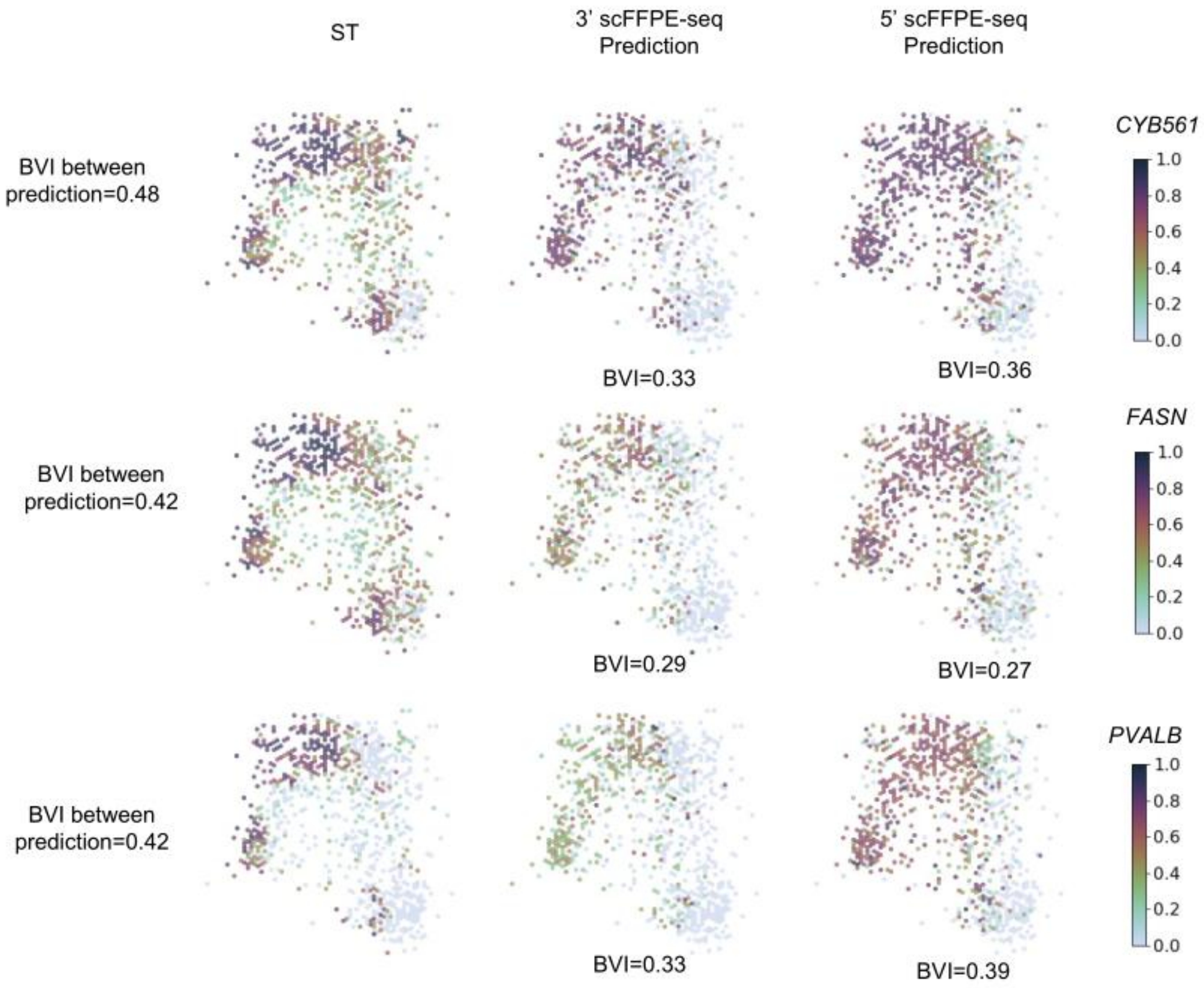


**Supplementary Figure 10. SC2Spa predicts spatial location of scFFPE-seq with high consistency to reference Visium data.** SC2Spa is trained using the Visium dataset for human breast cancer. The prediction results using 3' and 5' scFFPE-seq are shown for *CYB561*, *FASN* and *PVALB*. BVI is calculated between the reference ST and the prediction. The BVI between the two predictions (for 3' and 5' scFFPE-seq) is also shown to show the consistency.

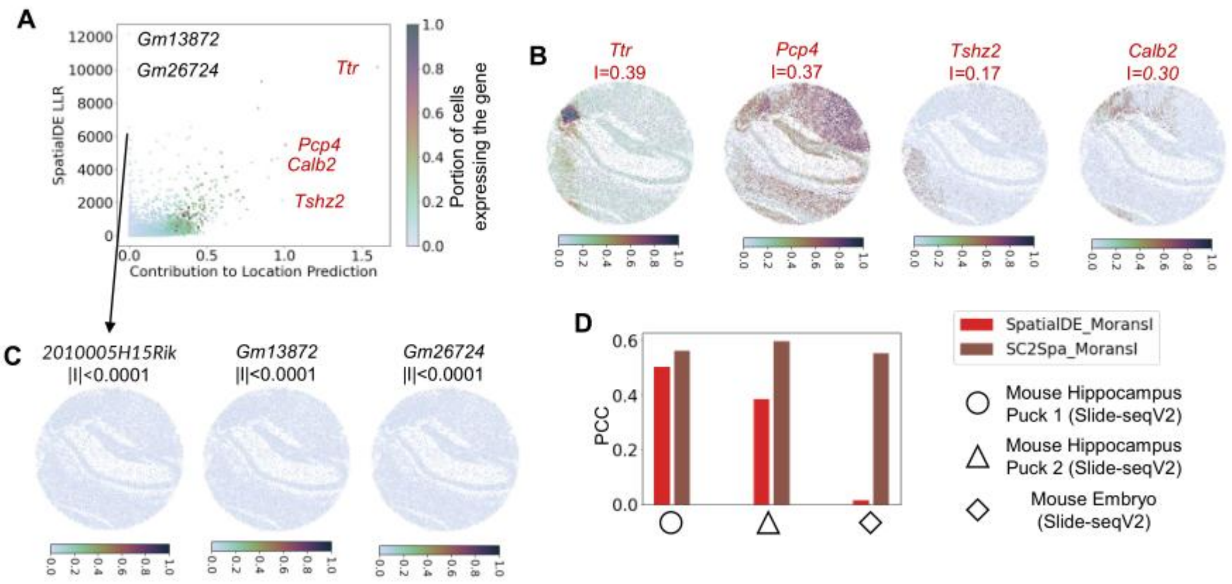


**Supplementary Figure 11. Assessing SVGs obtained by SC2Spa and SpatialDE. A.** The scatter plot for SC2Spa scores (contribution to location prediction) versus SpatialDE LLR values (PCC=0.562). The color gradient represents the percentage of cells where a gene is expressed. **B.** The SVGs commonly observed by SpatialDE and SC2Spa. **C.** The three genes with high SpatialDE LLR values but low SC2Spa scores. **D.** The PCC (Pearson's Correlation Coefficient) for SC2Spa and SpatialDE with Moran's I for three Slide-seqV2 datasets.

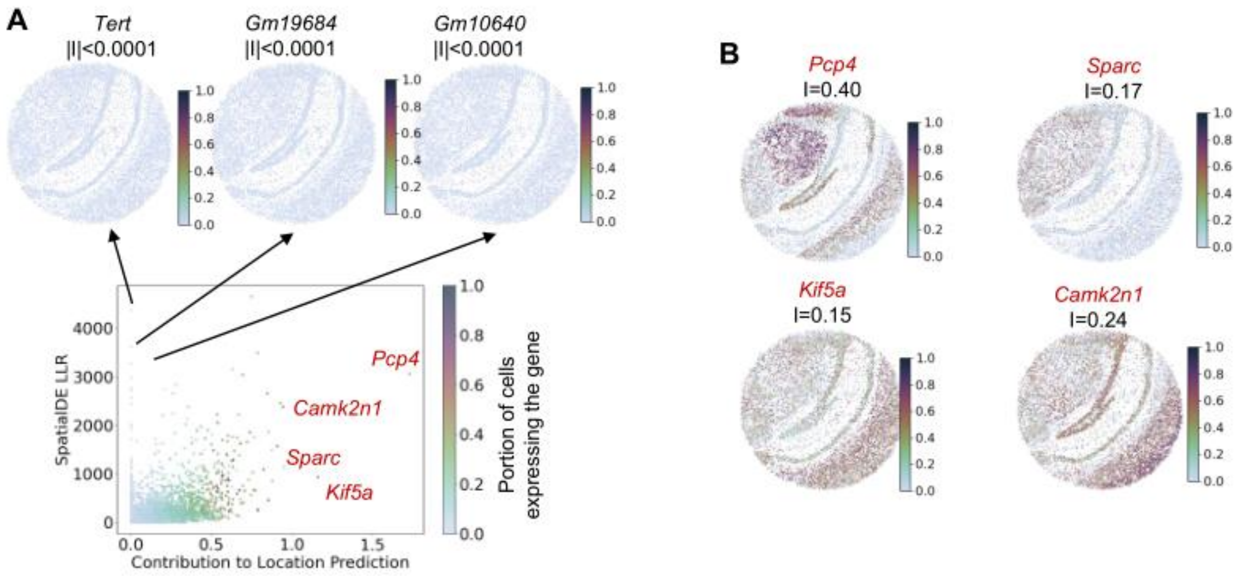


**Supplementary Figure 12. SC2Spa suggests SVGs from the mouse hippocampus Slide-seqV2 dataset (Puck_191204_01). A.** A scatter plot of SC2Spa score (contribution to location prediction) versus SpatialDE LLR value for all genes is shown. The color represents the proportion of cells that express a particular gene. *Tert* (I=-5.89e-5), *Gm19684* (I=-1.16e-4) and *Gm10640* (I=-3.15e-4) have high SpatialDE LLR scores but low SC2Spa scores. **B.** The top four genes identified by SC2Spa.

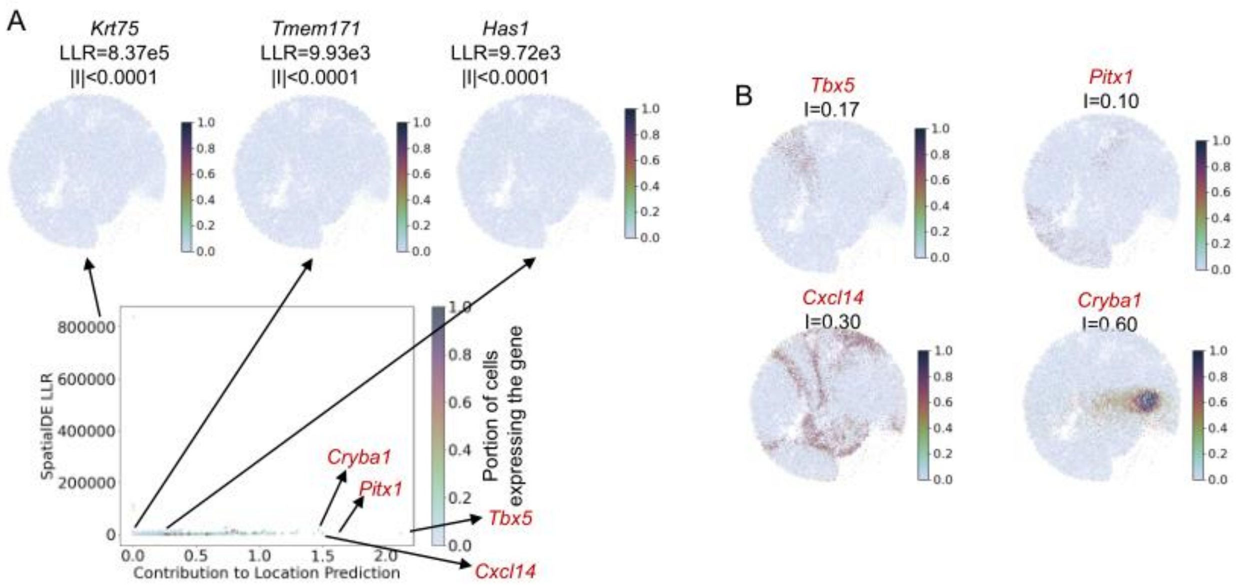


**Supplementary Figure 13. SC2Spa identified SVGs from the mouse embryo Slide-seqV2 data. A.** A scatter plot of SC2Spa score (contribution to location prediction) versus SpatialDE LLR value for all genes is presented. The hue indicates the proportion of cells that express a gene. *Krt75* (I=-6.90e-5), *Tmem171* (I=-6.94e-5) and *Has1* (I=-1.01e-4) have high SpatialDE LLR scores but low SC2Spa scores. **B.** The top four genes identified by SC2Spa.

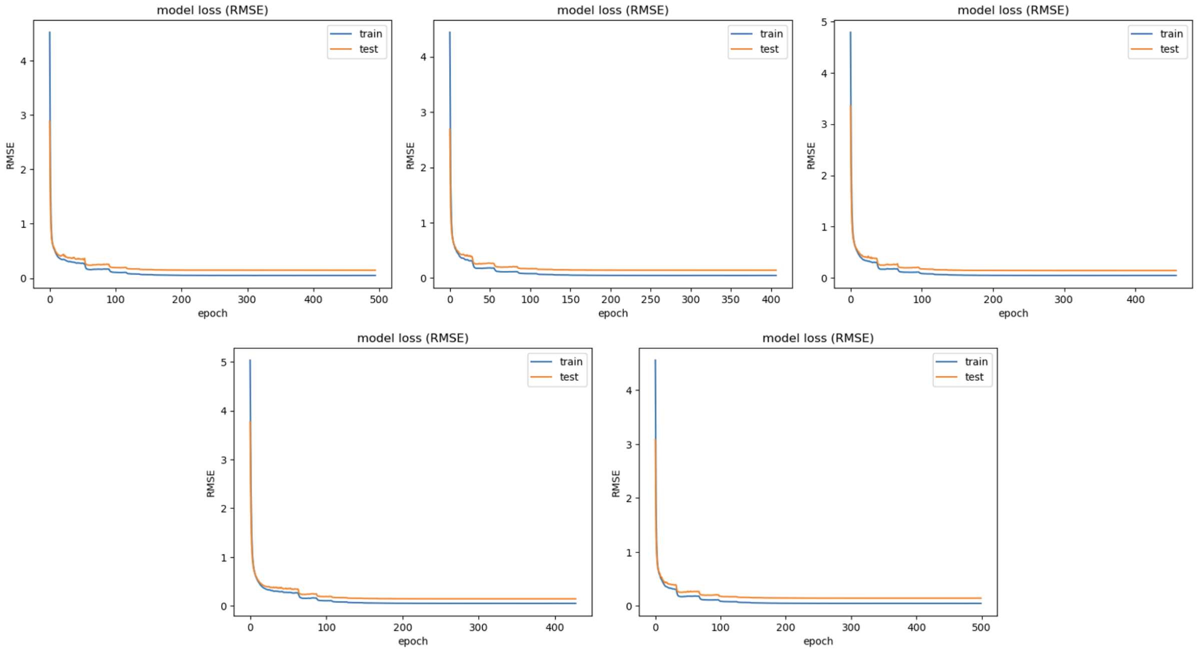


**Supplementary Figure 14. The RMSE value over training epoch.** RMSE histories of training and test data of the 5-fold cross-validation of Slide-seqV2 mouse hippocampus.

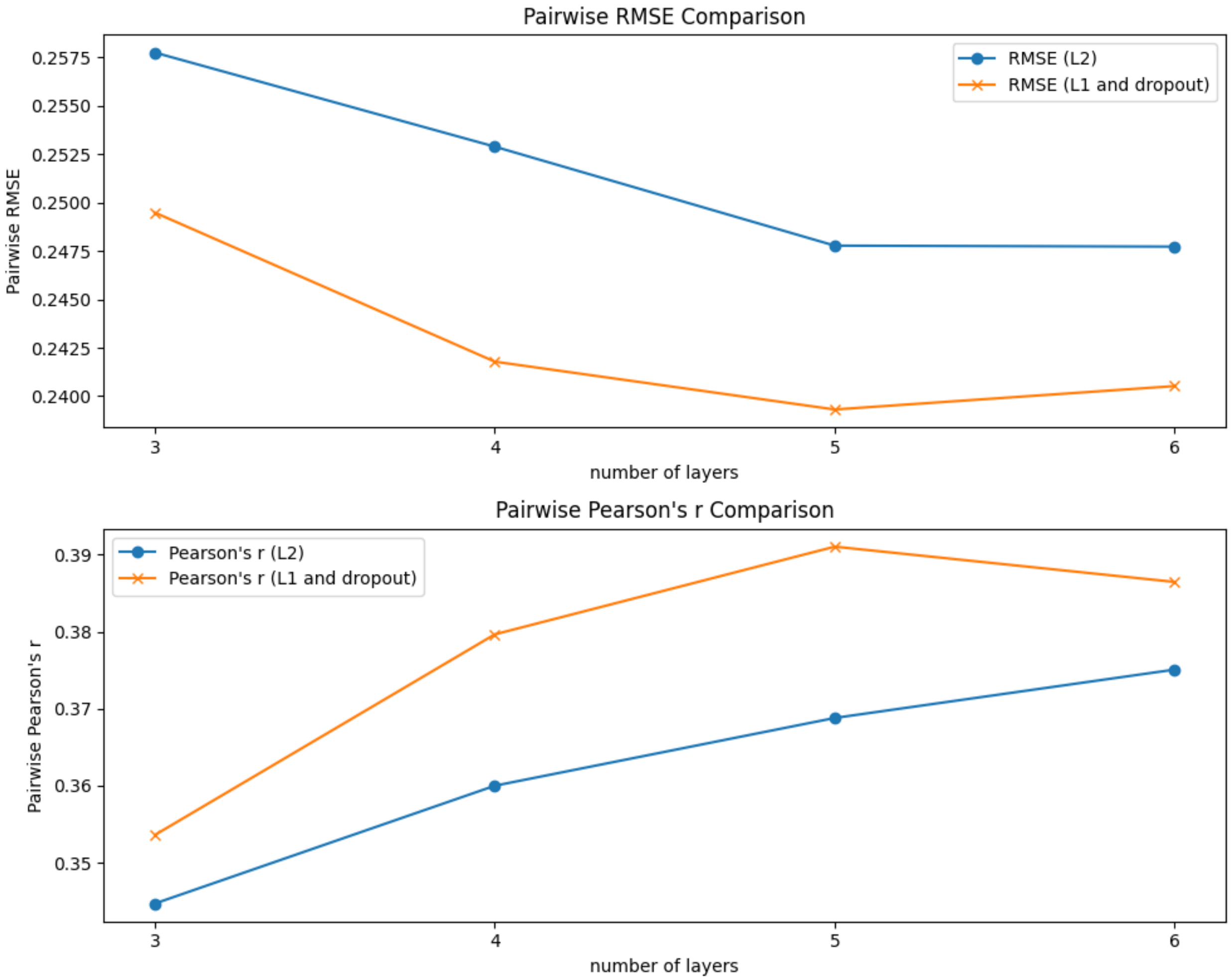


**Supplementary Figure 15. The effect of different factors for the performance of SC2Spa.** The pairwise Person's r and pairwise RMSE are plotted for SC2Spa using a varying number of layers in the model and L1 and L2 regularization on Slide-seqV2 mouse hippocampus data.

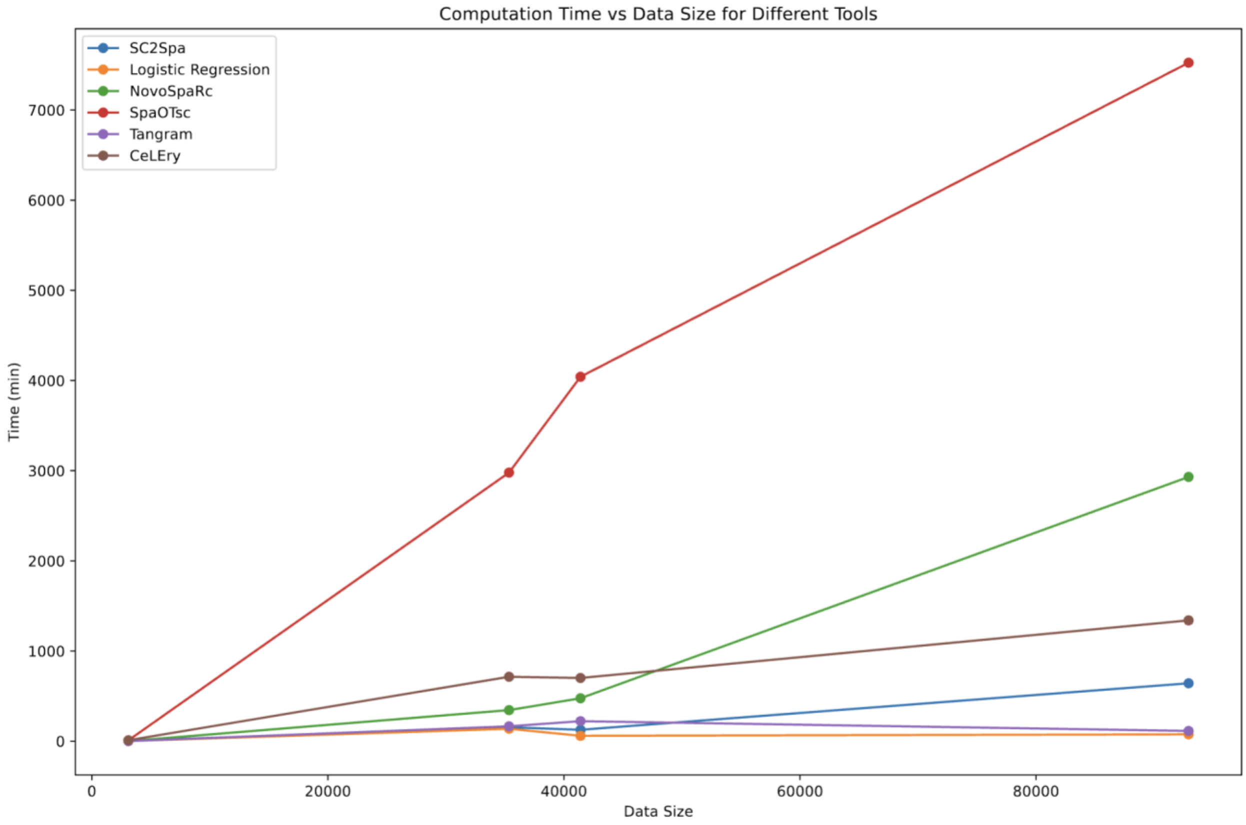


**Supplementary Figure 16. Comparison of running time across data size.** We compared the running time for each approach for the data size that we used for cross-validation.

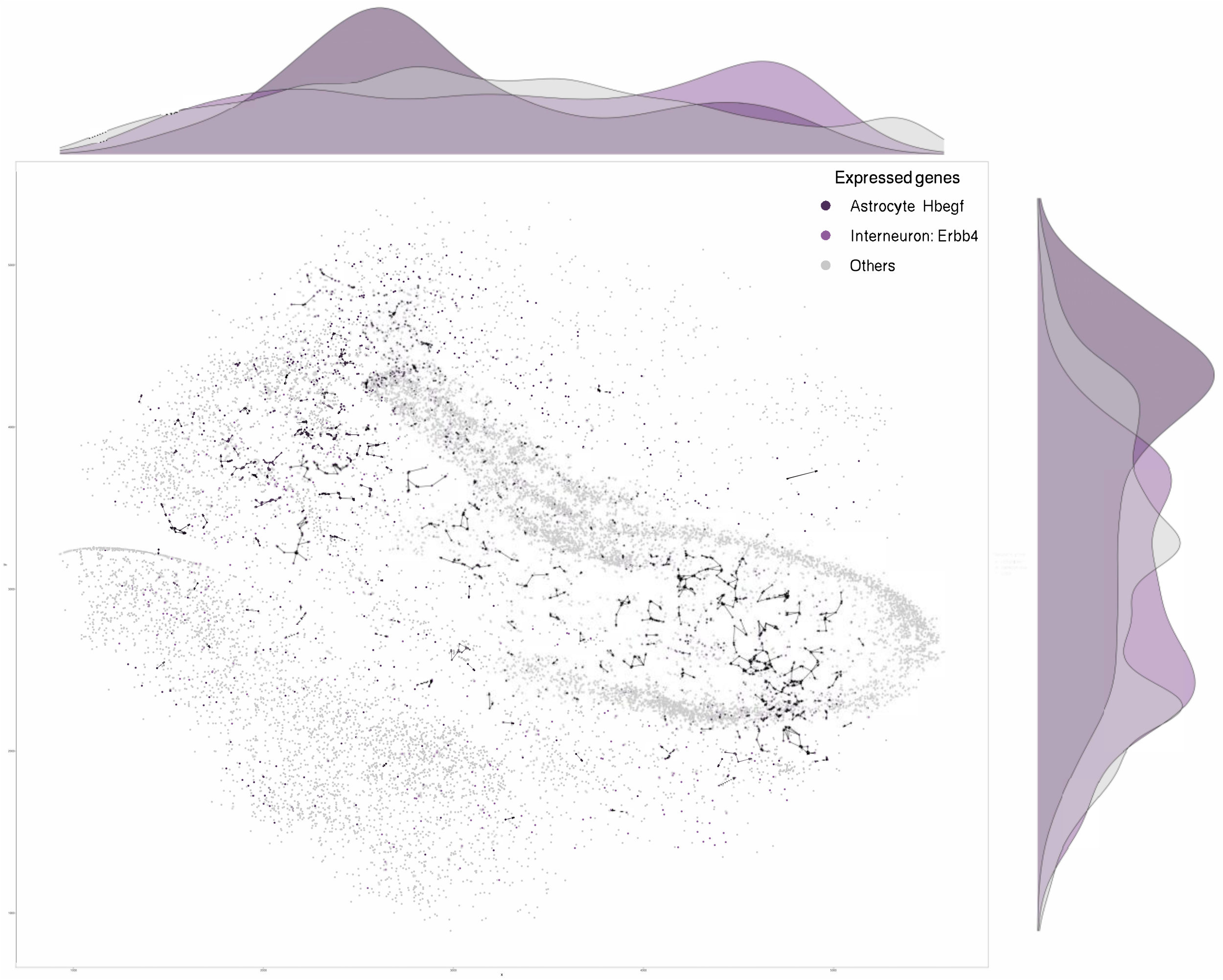


**Supplementary Figure 17. The spatial expression of Hbegf in interneuron and ErbB4 in Astrocyte.** The locations of interneurons and astrocytes were predicted using SC2Spa. The cell-cell communication was predicted using SpaTalk based on the results by SC2Spa.
